# Ciliogenic pancreatopathy reveals a link between ciliopathies and exocrine pancreatic disease

**DOI:** 10.64898/2026.01.27.701983

**Authors:** Memoona Rajput, Lydie Flasse, Esther Porée, Alice Serafin, Nicolas Papadopoulos, Younes Achouri, Axelle Loriot, Michaela Wilsch-Brauninger, Valentine Gillion, Nathalie Godefroid, Charlotte Bodson, Joel Moro, Leyre López-Muneta, Antonio Garcia de Herreros, Cécile Haumaitre, Frédéric Lemaigre, Meritxell Rovira, Amandine Viau, Anne Grapin-Botton, Sophie Saunier, Patrick Jacquemin, Isabelle Scheers

## Abstract

**Background:** While pancreatic cysts have been described in syndromic ciliopathies, the pancreas is not commonly recognized as a target organ. However, several ciliary gene knockout mouse models develop a pancreatic phenotype combining acinar atrophy and adipocyte accumulation, hereby called adipopancreatosis, suggesting a link between ciliary dysfunction and pancreatic disease.

**Objective:** We investigated whether mutations in ciliopathy-associated genes are linked to pancreatic dysfunction in humans.

**Design:** We analyzed a cohort of 341 patients with pediatric-onset pancreatic anomalies and characterized the pancreatic phenotype of new mouse models with conditional *Nphp3* inactivation or bearing *Nphp3* mutations recapitulating human mutations. In patients, pancreatic fat content was quantified using Dixon-MRI.

**Results:** Mutations in the cilium-related *HNF1B* and *NPHP3* were identified in patients presenting with both renal and pancreatic dysfunction. *Nphp3* mutant mice developed acinar atrophy, adipopancreatosis, and moderate inflammation. Adipocytes in the pancreas exhibited a white adipocyte-like profile and likely originated from mesothelial-derived fibroblasts. Reduced numbers and altered length of ductal cilia were monitored. Interestingly, secretory canaliculi, typically unnoticed structures found within and between acinar cells and connected to the acinar lumen, exhibited a microcystic morphology. Consistent with the mouse phenotype, Dixon-MRI revealed significantly increased pancreatic fat content in patients with *HNF1B* and *NPHP3* mutations.

**Conclusion:** We describe a previously unrecognized pancreatic manifestation of ciliopathies, which we name ciliogenic pancreatopathy. Patients with known ciliopathy-causing mutations should be evaluated for this pancreatic condition, particularly those with kidney disease, as concomitant exocrine pancreatic insufficiency may further compromise renal function or the outcome of kidney graft.

**What is already known on this topic:**

- Ciliopathies, resulting from defects in primary cilia, are genetic disorders primarily affecting the kidney and liver.
- Pancreatic cysts have been sporadically reported in syndromic ciliopathies.
- The pancreas is not currently recognized as a major target organ of ciliary dysfunction.
- A clear link between ciliary gene mutations and pancreatic anomalies is still unknown.
- Animal studies have suggested a possible association between ciliary dysfunction and pancreatic anomalies.

**What this study adds:**

- Identifies HNF1B and NPHP3 mutations as genetic causes of a pancreatic phenotype characterized by acinar atrophy and adipose replacement (adipopancreatosis).
- Demonstrates the presence of defective ductal cilia and moderate inflammation in the pancreas of *Nphp3* mutant mice.
- Reveals that secretory canaliculi in the exocrine pancreas of *Nphp3* mutant mice acquire a microcystic morphology.
- Shows that patients with HNF1B or NPHP3 mutations have significantly increased pancreatic fat content by Dixon-MRI.
- Defines a new disease entity, ciliogenic pancreatopathy, as a pancreatic manifestation of ciliopathies.

**How this study might affect research, practice or policy:**

- Establishes the pancreas as a novel and clinically relevant target of ciliopathies.
- Expands the phenotypic spectrum of HNF1B- and NPHP3-related diseases to include exocrine pancreatic dysfunction.
- Suggests that patients with ciliopathy-causing mutations should be evaluated for exocrine pancreatic insufficiency.
- Highlights the need to consider pancreatic function monitoring in kidney disease and transplant settings.
- Opens new research avenues into the role of primary cilia in pancreatic homeostasis and disease.

## Introduction

A number of pathological conditions involving the exocrine pancreas remain poorly defined or not well understood. These include cases of idiopathic pancreatic dysfunction, in which patients experience exocrine pancreatic insufficiency leading to maldigestion and diarrhea, or conditions that mimic pancreatitis but do not exhibit all the typical symptoms or associated risk factors. Their diagnosis can be challenging due to clinical variability, rarity, or the absence of precise diagnostic criteria^1^.

In terms of pancreatic pathology, the presence of cysts has been associated with certain ciliopathies, such as polycystic kidney disease and Von Hippel-Lindau syndrome^2, 3^. Ciliopathies are a group of rare genetic disorders caused by abnormalities of the motile or primary cilia. Unlike motile cilia, primary cilia are immotile structures present on the plasma membrane of many cell types. They play important roles in cell signaling and organ development during embryogenesis. As a result, dysfunction of the cilia can lead to various complications in organs such as the kidneys, central nervous system, and liver^4^. Aside from the cysts mentioned above, the pancreas is not usually recognized as a primary target organ in ciliopathies.

In the pancreas, primary cilia are absent from acinar cells but are present on both ductal and endocrine cells. Interestingly, multiple mouse models with inactivation of ciliary or ciliogenic genes consistently develop pancreatic inflammation and adipopancreatosis, accompanied by acinar atrophy. These include models with duct-specific postnatal deletion of *Hnf6*, *Stk11/Lkb1*, and *Hnf1B*, all of which impair ciliary function^5–6^, as well as knockouts for *Ift88* and *Kif3a*^7–8^, among others. Adipopancreatosis (from adipo- for adipose tissue, pancreat- for pancreas, and -osis for a pathological state) refers to the accumulation of adipocytes within the pancreatic parenchyma, accompanied by tissue atrophy. This suggests a potential causal link between cilium dysfunction and these specific pancreatic anomalies, although further evidence is needed both in animal models and even more so in humans.

In the present study, we identified mutations in the *HNF1B* and *Nephrocystin 3* (*NPHP3*) genes in a cohort of patients, including children and adolescents, who presented with pancreatic dysfunction and features of ciliopathies. Characterization of several mouse models with ciliary dysfunction then led us to hypothesize that impaired homeostatic interactions between pancreatic parenchymal cell types lead to the development of adipopancreatosis and acinar atrophy. Re-evaluation of the cohort using medical imaging revealed an increased pancreatic fat content (PFC) in these patients. Collectively, these results support the existence of a previously unrecognized pancreatic disease, which we name ciliogenic pancreatopathy.

## Materials and Methods

Detailed descriptions of all materials and experimental methods can be found in online supplementary methods.

## Results

### Patients with *HNF1B* and *NPHP3* mutations exhibit ciliopathic features associated with pancreatic abnormalities

As part of the European Reference Network for rare Inherited and Congenital (digestive and gastrointestinal) Anomalies (ERNica), the CuSL expert center currently follows a cohort of 341 patients with pediatric-onset pancreatic anomalies, established since 2016. Routine biochemistry, imaging, and genetic testing identified common causes of pancreatic disease in most cases. However, 30 patients (8.8%) remained without a clear diagnosis and were classified as having idiopathic pancreatic disease.

Interestingly, 6 of these idiopathic patients were already receiving nephrological care for kidney problems associated with ciliopathy (Figure 1A). Genetic testing revealed that 4 of these patients carried a mutation in the *HNF1B* gene (HNF1B_1-4), while the other 2 had biallelic mutations in the *NPHP3* gene (NPHP3_1-2) (Figure 1A). *HNF1B* is known to control ciliary gene expression^6–8^ and its deficiency has been linked to ciliary defects in human cholangiocytes^8, 30, 31^. Furthermore, *NPHP3* is frequently mutated in patients with nephronophthisis (NPH), an autosomal recessive ciliopathy affecting the kidney^32^. From this result, we identified 24 additional patients followed in nephrology and carrying *HNF1B* or *NPHP3* mutations, namely 21 HNF1B patients (HNF1B_ 5-25) and 3 NPHP3 patients (NPHP3_3-5), and explored their pancreatic function. Nine HNF1B patients (HNF1B_5-13) presented with pancreatic abnormalities while the other HNF1B patients did not have apparent pancreatic deficiencies (Figure 1A).

**Figure 1.**
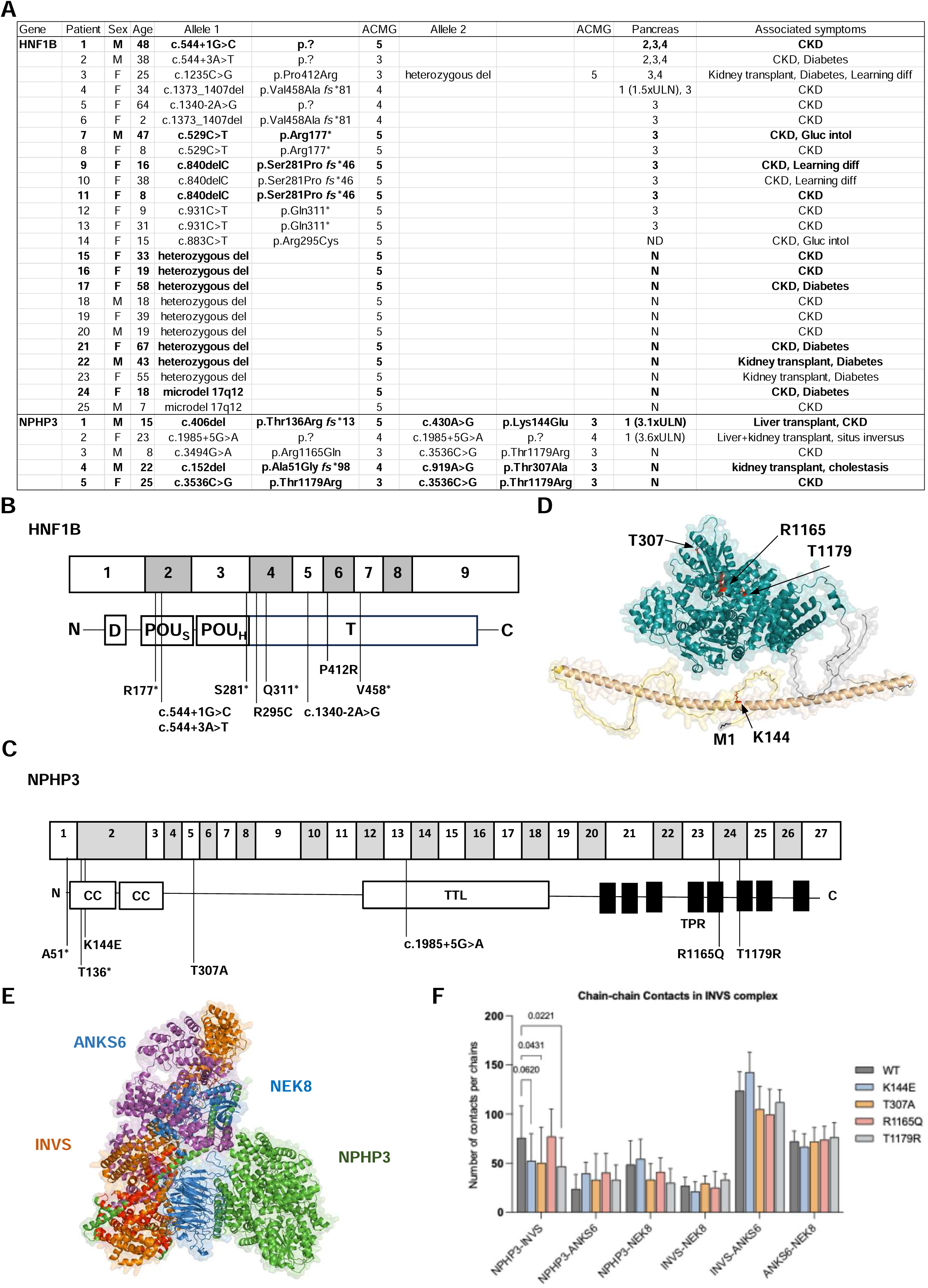
Ciliopathic features and pancreatic abnormalities are associated with mutations in the transcription factor HNF1B and the ciliary protein NPHP3. (A) Genotype and clinical characteristics of the CuSL patient cohort with *NPHP3* and *HNF1B* mutations. Patients in bold underwent MRI to determine PFC. Sex is indicated as F: female, or M: male. Age is indicated in years. ACMG: variant classification according to the American College of Medical Genetics, where (3) corresponds to a mutation classified as variant of unknown pathogenic significance, (4) likely pathogenic, and (5) pathogenic. The “Pancreas” column indicates pancreatic anomalies observed in patients: (1) raised lipase, (2) exocrine pancreatic insufficiency, (3) abnormal imaging findings, (4) history of acute, acute recurrent, or chronic pancreatitis; xULN, fold above the upper limit of normal; Gluc intol, glucose intolerance, N: no pancreatic anomaly, ND: no data. The “Associated symptoms” column lists additional clinical features: CKD: chronic kidney disease; del: deletion; Learning diff: learning difficulties. (B) Schematic representation of the *HNF1B* gene and protein structure with mutations reported in patients. D, dimerization domain; POU_S_ and POU_H_, DNA binding domains of HNF1B Pit-Oct-Unc (POU)-specific domain and POU homeodomain; T, transactivation domain. (C) Schematic representation of the *NPHP3* gene and protein structure with mutations reported in patients. CC, coiled-coil domain; TTL, tubulin tyrosine ligase domain; TPR, tetratricopeptide-repeat domain. (D) Structure prediction of NPHP3 performed with AlphaFold 3.0. The unstructured N-terminal domain is shown in yellow, the α-helical region in wheat, connecting linkers in grey, and the globular domain in deep blue. (E) Structure prediction of INVSc using AlphaFold 3.0. ANKS6 is shown in purple, INVS in orange, NEK8 in dark blue, and NPHP3 in green. Contact residues between INVS and NPHP3 (distance < 6Ä) are shown in red. (F) Number of chain-chain contacts for indicated pairs. Contact pairs are defined as residues whose carbon ß (or carbon α for Gly) have a radial distance < 6Ä. Data represent mean + SD of 5 AlphaFold 3.0 models per condition. Statistics: two-ways ANOVA with Fisher’s LSD test (uncorrected).

Among the 25 HNF1B patients, 12 carried point mutations and 13 had gene deletions, including 2 with a larger 17q12 microdeletion. The median age of HNF1B patients was 31 years (range: 2-67 years), and 18 were women. Three patients had chronic pancreatitis (CP), one of them being a heavy alcohol consumer (HNF1B_1), and one had chronically raised lipase levels (HNF1B_4). Among the NPHP3 patients, 2 carried homozygous point mutations, and 3 were compound heterozygotes (Figure 1A). Their median age was 18.6 years (range: 8-25 years), and 2 were women. Two patients (NPHP3_1-2) had chronically raised lipase levels. HNF1B and NPHP3 mutations were often found in conserved protein domains (Figure 1B-C). Extensive genetic, metabolic and imaging evaluations have not identified any well-defined cause of exocrine pancreatic diseases. Demographic and clinical information are provided in Figure 1A. Together, these results suggest that a subset of idiopathic pancreatic conditions may be linked to underlying ciliary dysfunction.

While HNF1B is an important transcription factor for ciliary gene expression^6, 23^, NPHP3 is localized in the cilium. In fact, NPHP3 is a component of INVSc, a ciliary tetrameric assembly also comprising INVS, NEK8, and ANKS6^33^. Therefore, we assessed how the distinct NPHP3 missense mutations identified in our patient cohort may alter the protein’s structure and its interactions with the proteins of INVSc. To gain structural insight, NPHP3 alone or as part of INVSc was modeled using AlphaFold 3.0^13^. The predicted structure of NPHP3 alone features an unstructured N-terminal region, followed by an extended α-helix linked through a short, disordered segment to a globular domain. The latter encompasses most of the protein, including the C-terminal tetratricopeptide repeats (TPR), whereas the C-terminal part ends in a short unstructured tail (Figure 1C-D).

Interestingly, when interacting with the INVSc members, the NPHP3 N-terminus was predicted to form α-helices, rather than random coils. Notably, the simulations also indicated a direct interaction between the N-terminal region of NPHP3 and the C-terminal domain of INVS, a feature not previously described. The NPHP3 globular domain primarily interacted with NEK8, while INVS was predominantly associated with ANKS6 (Figure 1E). In agreement with experimental data, all INVSc models consistently positioned the NEK8–ANKS6 dimer at the core of the complex^33^.

NPHP3 mutants, except R1165Q, were predicted to decrease the number of contacts (defined as residues closer than 6 Ä) between the NPHP3 N-terminus and the INVS C-terminus (Figure 1F). In the case of T307A, and for T1179R in some models, this loss of contacts coincided with a topological rearrangement of the complex (Supplementary Figure 1A). Taken together, this *in silico* analysis suggests that the missense mutations weaken the NPHP3–INVS interaction, potentially compromising the integrity of INVSc. Supporting this, DynaMut analysis revealed that all substitutions, except the K144E substitution, destabilize the tetrameric complex (Supplementary Figure 1B), potentially explaining their deleterious effects.

### A patient-specific mouse model carrying *NPHP3* mutations replicates the kidney manifestations of nephronophthisis type 3

Mouse models with *Hnf1B* mutations exhibit pancreatic acinar loss and adipopancreatosis^8, 23^, prompting us to investigate whether *Nphp3* mutations result in a comparable phenotype. To address this, we generated a CRISPR/Cas9-engineered mouse model reproducing the two *NPHP3* mutations identified in the first patient (NPHP3_1) using CRISPR/Cas9 technology. Thus, we inserted the mutations c.655A>G and c.631del (from the mouse *Nphp3* cDNA sequence NM_028721), referred to as mut1 and mut2 respectively, into the mouse *Nphp3* gene to obtain the so-called *Nphp3*^mut1/mut2^ patient-specific mouse model (Supplementary Figure 2A-B). Mut1 is expected to generate a hypomorphic allele, while mut2 is expected to result in a null allele.

Before studying the pancreatic phenotype of these mice, we sought to validate this model by showing that it recapitulated the main renal characteristics of nephronophthisis (NPH) type 3. A slight reduction in *Nphp3* mRNA expression was observed in kidneys from *Nphp3*^mut1/mut2^ animals compared to controls (Supplementary Figure 3A). Analysis of the kidneys from *Nphp3*^mut1/mut2^ mice revealed the progressive development of tubular cysts. Cysts were evident as early as 2 months after birth, and cyst size and number progressively increased with age (Figure 2A). Tubular cysts arise from the thick ascending limb of the loop of Henle (TAL) and the distal part of the nephron, mainly from the connecting tubules (CNT) and the collecting ducts (CD) (Figure 2B-C). While no tubular cysts were observed in the proximal part of the nephron, glomerulocysts were often observed. These histologic features resemble those reported in *pcy* mice^34^, which carry a hypomorphic *Nphp3* allele^35^. Over time, *Nphp3*^mut1/mut2^ kidneys showed reduced *Uromodulin* and *Calbindin 1* mRNA expression compared to controls, suggesting dedifferentiation of the thick ascending limb of the loop of Henle and distal tubules, respectively (Figure 2 D-E). At late stage, some tubular atrophy and thickening of tubular basement membrane were associated with increased expression of known tubular injury markers (*Havcr1*, *Lcn2*) in *Nphp3*^mut1/mut2^ kidneys in comparison to controls (Figure 2F-G). Progressive diffuse interstitial fibrosis was found in cystic *Nphp3*^mut1/mut2^ kidneys in comparison to controls, with deposition of collagen fibers from 12 months onward (Figure 2H). Despite this cystic phenotype, *Nphp3*^mut1/mut2^ animals did not present kidney failure (Supplementary Figure 3B).

**Figure 2.**
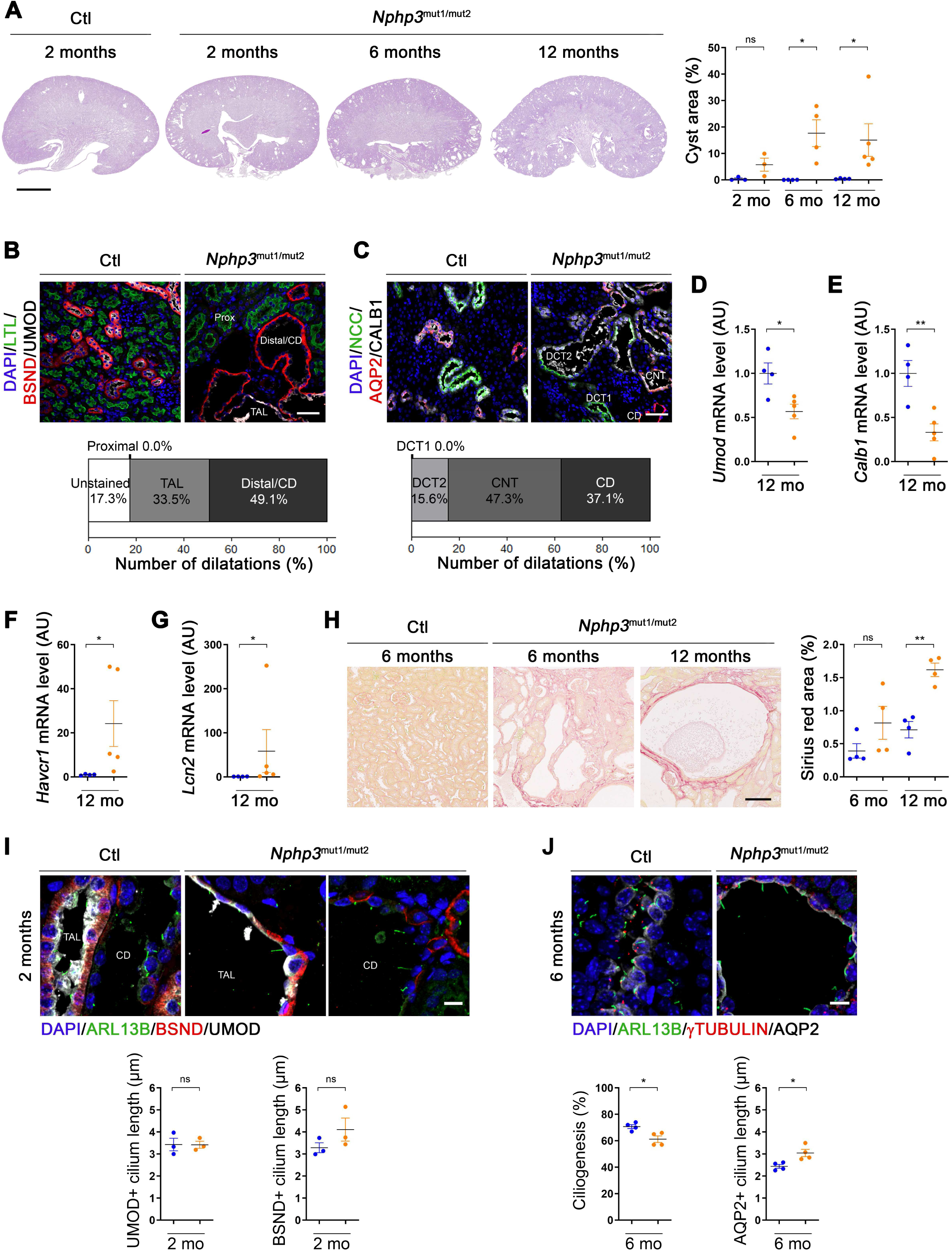
*Nphp3* mutations induce cystic growth, tubular injury, interstitial fibrosis, and primary cilium defects in the kidneys. **(A)** Representative PAS staining of kidney sections and quantification of cyst area fraction from control and *Nphp3*^mut1/mut2^ mice at the indicated age. Scale bar: 2.5 mm. **(B)** Immunolabeling and quantification of kidney sections from control and *Nphp3*^mut1/mut2^ mice at 2 months with LTL (for proximal tubules, green), UMOD (for TAL, grey) and BSND (for TAL, distal tubules and CD, red) antibodies and DAPI (nuclei, blue). Scale bar: 50 µm. n=3. **(C)** Immunolabeling and quantification of kidney sections from control and *Nphp3*^mut1/mut2^ mice at 2 months with NCC (for DCT1 and DCT2, green), CALB1 (for DCT2 and CNT, grey), AQP2 (for CNT and CD, red) antibodies and DAPI (nuclei, blue). Scale bar: 50 µm. n=3. **(D-G)** *Umod* (D), *Calb1* (E), *Havcr1* (F) and *Lcn2* (G) mRNA content evaluated by qRT-PCR in controls and *Nphp3*^mut1/mut2^ mice kidneys at 12 months. **(H)** Representative Sirius red-stained kidney sections and quantification of interstitial fibrosis from control and *Nphp3*^mut1/mut2^ mice at the indicated age. Scale bar: 100 µm. **(I)** Immunolabeling and quantification of kidney sections from control and *Nphp3*^mut1/mut2^ mice at 2 months with ARL13B (for primary cilia, green), UMOD (for TAL, grey) and BSND (for TAL, distal and CD, red) antibodies and DAPI (nuclei, blue). Scale bar: 10 µm. n=50 cilia counted per mouse. **(J)** Immunolabeling and quantification of kidney sections from control and *Nphp3*^mut1/mut2^ mice at 2 months with ARL13B (for primary cilia, green), γTUBULIN (for basal body, red) and AQP2 (for CD, grey) and DAPI (nuclei, blue). Scale bar: 10 µm. n=50 cilia counted per mouse for cilium length quantification and n=100 cilia counted per mouse for ciliation assessment. **(A-J)** Each dot corresponds to one mouse, with controls shown in blue and *Nphp3*^mut1/mut2^ mice in orange. * P < 0.05, ** P < 0.01. AU: arbitrary unit. TAL: Thick Ascending Limb of the Loop of Henle, DCT: Distal Convoluted Tubule, CNT: Connecting Tubule, CD: Collecting Duct. Blue dot, Control; orange dot, *Nphp3*^mut1/mut2^.

As NPHP3 encodes a protein that localizes to the primary cilium and contributes to its biogenesis^36, 37^, we next investigated cilium morphology and number. At 2 months, cystic tubules in *Nphp3*^mut1/mut2^ kidneys presented with ARL13B-positive primary cilia of normal length in both the thick ascending limb of the loop of Henle and distal tubules (Figure 2I). However, by 6 months, while the percentage of ciliated cells in the cysts slightly decreased, primary cilia were elongated in cystic epithelial cells of *Nphp3*^mut1/mut2^ kidneys in comparison to controls (Figure 2J and Supplementary Figure 3C). Overall, the close similarities between the renal abnormalities in *Nphp3*^mut1/mut2^ mice and those observed in the kidneys from NPHP3 patients^38^ support the validity of this model for studying NPH type 3.

### The pancreas of *Nphp3* patient-specific model is characterized by progressive acinar atrophy and adipopancreatosis

To characterize the pancreatic phenotype of *Nphp3*^mut1/mut2^ mice, we first performed histological and immunohistochemical analyses of pancreata from 2- to 8-month-old mice (Figure 3A). From 6 months, we observed a progressive accumulation of adipocytes within the pancreatic parenchyma of *Nphp3*^mut1/mut2^ mice, which increased at 8 months. Notably, this adipocyte infiltration was not associated with fibrosis, and no lymphocytic infiltration was detected at any time point examined (Supplementary Figure 4). However, by 8 months of age, a significant increase in F4/80-positive cells was observed in the pancreas of *Nphp3*^mut1/mut2^ mice compared to controls. In contrast to the few round-shaped peritoneal macrophages occasionally observed in the pancreatic mesothelium of control pancreas, the F4/80-positive cells in the mutant pancreas exhibited an elongated morphology and were frequently located at the periphery of pancreatic lobules (Figure 3A-B).

**Figure 3.**
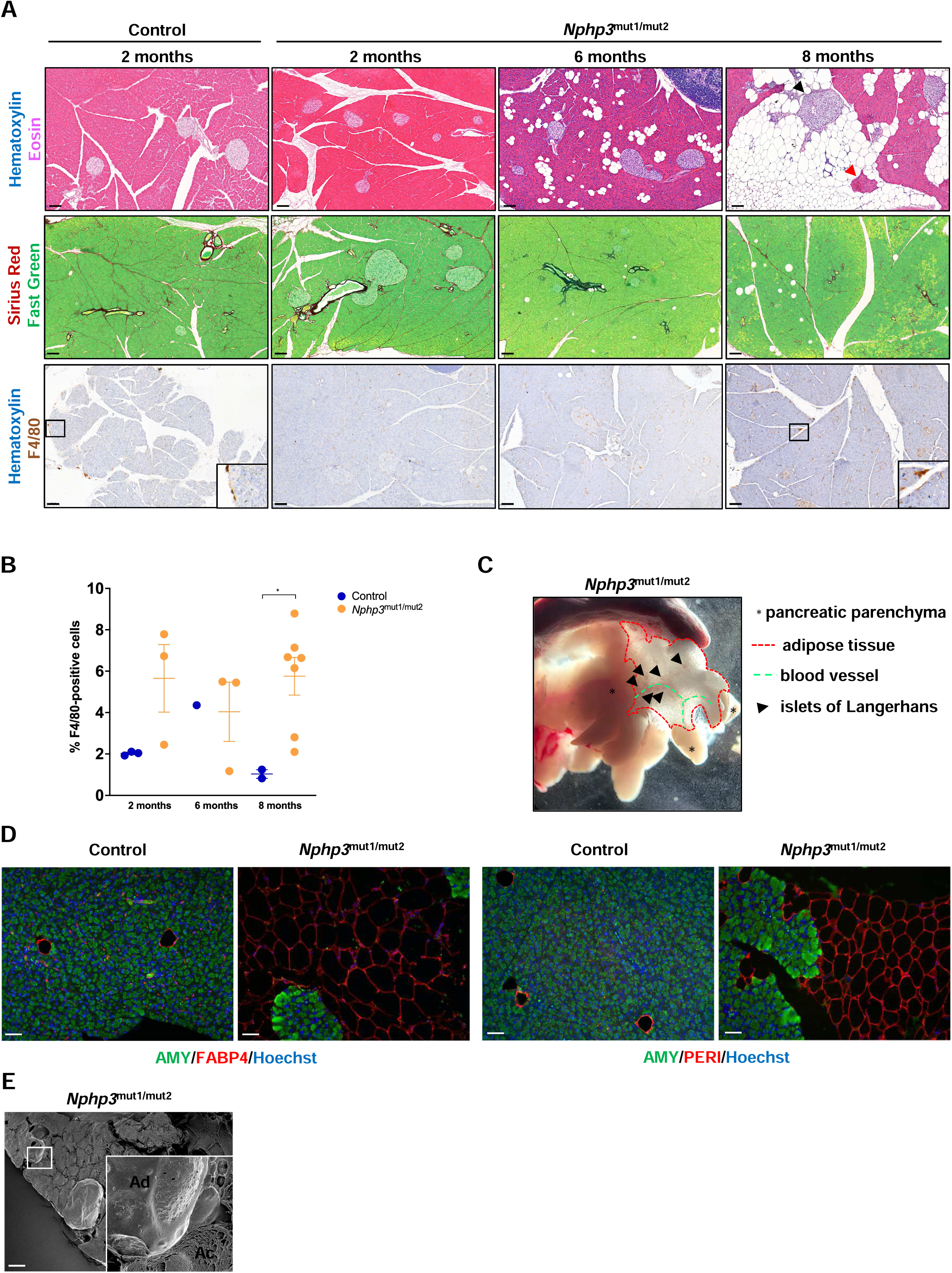
*Nphp3*^mut1/mut2^ mice exhibit progressive lipomatosis, leading to acinar atrophy. **(A)** Hematoxylin/Eosin, Sirius Red/Fast Green, and Hematoxylin/F4/80 staining of pancreas sections from control and *Nphp3*^mut1/mut2^ mice at different time points. Scale bar: 100 μm. n=3-5. **(B)** Quantification of the percentage of F4/80-positive cells in the pancreas of Control and *Nphp3*^mut1/mut2^ mice. **(C)** Picture of a dissected pancreas from a 10-month-old *Nphp3*^mut1/mut2^ mouse. **(D)** Immunolabeling of pancreas sections from control and *Nphp3*^mut1/mut2^ mice at 8 months with amylase (for acinar cells), FABP4 and perilipin 1 (PERI. For adipocytes) antibodies and Hoechst (nuclei, blue). Non-specific labeling of individual cells, probably red blood cells, is observed with the FABP4 antibody on the Control section. Scale bar: 50 μm. FABP4: Control n= 3 and *Nphp3*^mut1/mut2^ n=3; Perilipin 1: Control n=3 and *Nphp3*^mut1/mut2^ n=3. **(E)** Scanning electron microscopy of a pancreas section from a *Nphp3*^mut1/mut2^ mouse at 8 months. Scale bar: 20 μm. Ad, adipocyte; Ac, acinar cell.

Adipopancreatosis showed a highly variable penetrance between lobes. In some *Nphp3*^mut1/mut2^ mice, entire lobes of the pancreas appeared to be replaced by adipocytes (Figure 3C). Within these adipocyte-rich regions, residual islets of Langerhans and occasional acinar structures remained detectable (Figure 3A and 3C).

These adipocytes expressed the classical adipocyte markers FABP4 and perilipin1 (PERI) (Figure 3D) and were found in direct contact with acinar cells (Figure 3E). Together, these observations indicate that the pancreas of *Nphp3*^mut1/mut2^ mice develops progressive acinar atrophy associated with adipopancreatosis, in the context of a mild F4/80-positive cell infiltration.

### Ductal deletion of *Nphp3* also leads to progressive acinar atrophy and adipopancreatosis

Since ductal cells are the only ciliated cell type of exocrine pancreas and no ductal abnormalities are observed in the *Nphp3*^mut1/mut2^ pancreas, we next investigated whether conditional deletion of *Nphp3* in this cell type might contribute to the pancreatic phenotype. To address this question, we generated a conditional knockout mouse model called *Nphp3*^f/f^ (Supplementary Figure 2C). These mice were then crossed with *Sox9Cre^ER^* mice to specifically inactivate *Nphp3* in ductal cells.

First, we investigated whether the specificity of the *Sox9Cre^ER^* model was maintained in the *Nphp3*^f/f^ background. Accordingly, we used *Rosa^YFP^* reporter mice and confirmed that Cre^ER^ activity was strictly restricted to ductal cells, as demonstrated by co-labeling of GFP and SOX9 both in *Sox9Cre^ER^ Rosa^YFP^* and *Sox9Cre^ER^ Nphp3*^f/f^ *Rosa^YFP^* pancreata. We also observed high recombination efficiency at the *Rosa* locus within the ductal compartment (Figure 4A). Next, we performed histological and immunohistochemical analyses identical to those performed on the *Nphp3* patient-specific model. Ductal hyperplasia was observed (Figure 4B). As in the patient-specific model, we observed the presence of adipopancreatosis. In contrast, mild fibrosis was present in the *Sox9Cre^ER^ Nphp3*^f/f^ pancreas (Figure 4B). CD3-positive lymphocytic infiltrates were observed in some lobes, accompanied by a significant increase in

**Figure 4.**
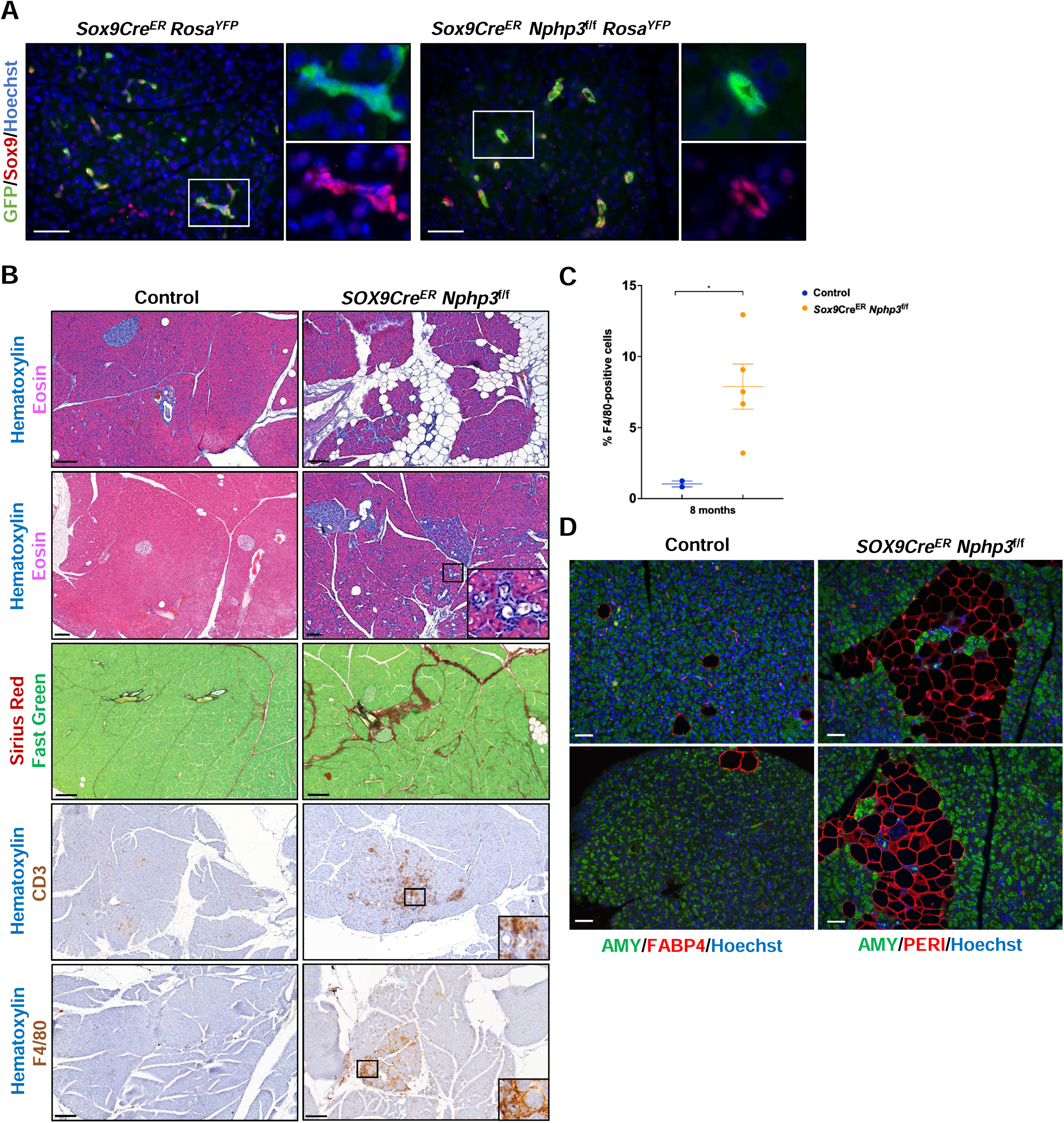
Ductal inactivation of *Nphp3* leads to lipomatosis, and mild fibrosis and inflammation. **(A)** Immunolabeling of pancreas sections from *Sox9Cre^ER^ Rosa^YFP^* and *Sox9Cre^ER^ Nphp3*^f/f^ *Rosa^YFP^* mice at 8 months with SOX9 (for ductal cells) and GFP (for ductal cell tracing) antibodies, and Hoechst (nuclei, blue). **(B)** Hematoxylin/Eosin, Sirius Red/Fast Green, Hematoxylin/CD3, and Hematoxylin/F4/80 staining of pancreas sections from control and *Nphp3*^mut1/mut2^ mice at different time points. Scale bar: 100 μm. **(C)** Quantification of the percentage of F4/80-positive cells in the pancreas of Control and *Sox9Cre^ER^ Nphp3*^f/f^ mice. **(D)** Immunolabeling of pancreas sections from control and *Sox9Cre^ER^ Nphp3*^f/f^ mice at 8 months with amylase (for acinar cells), FABP4 and perilipin 1 (PERI. For adipocytes) antibodies and Hoechst (nuclei, blue). n=4-6. A and E, scale bar: 50 μm.

F4/80-positive macrophages within the affected areas (Figure 4B-C). As expected, adipocytes present in regions of adipopancreatosis expressed FABP4 and perilipin1 (Figure 4D). These results show that acinar atrophy and adipopancreatosis develop following *Nphp3* inactivation in ductal cells. Moreover, they reveal that a complete loss of NPHP3 function in this cell type induces a more pronounced immune infiltration compared to the phenotype resulting from hypomorphic NPHP3 activity.

### Patient-derived mutations and ductal inactivation of *Nphp3* affect the primary cilium

Given that NPHP3 encodes a protein localized to the primary cilium, we investigated whether patient-derived *Nphp3* mutations or ductal-specific *Nphp3* deletion impact ciliary structure. To assess ciliary integrity, we performed immunofluorescence labeling for ARL13B, in combination with CK19 to identify ductal cells on pancreas sections. In *Nphp3*^mut1/mut2^ pancreas, primary cilia in ductal cells were significantly elongated compared to controls. In contrast, primary cilia appeared significantly shorter in Sox9Cre^ER^ *Nphp3*^f/f^ pancreas (Figure 5A-B).

**Figure 5.**
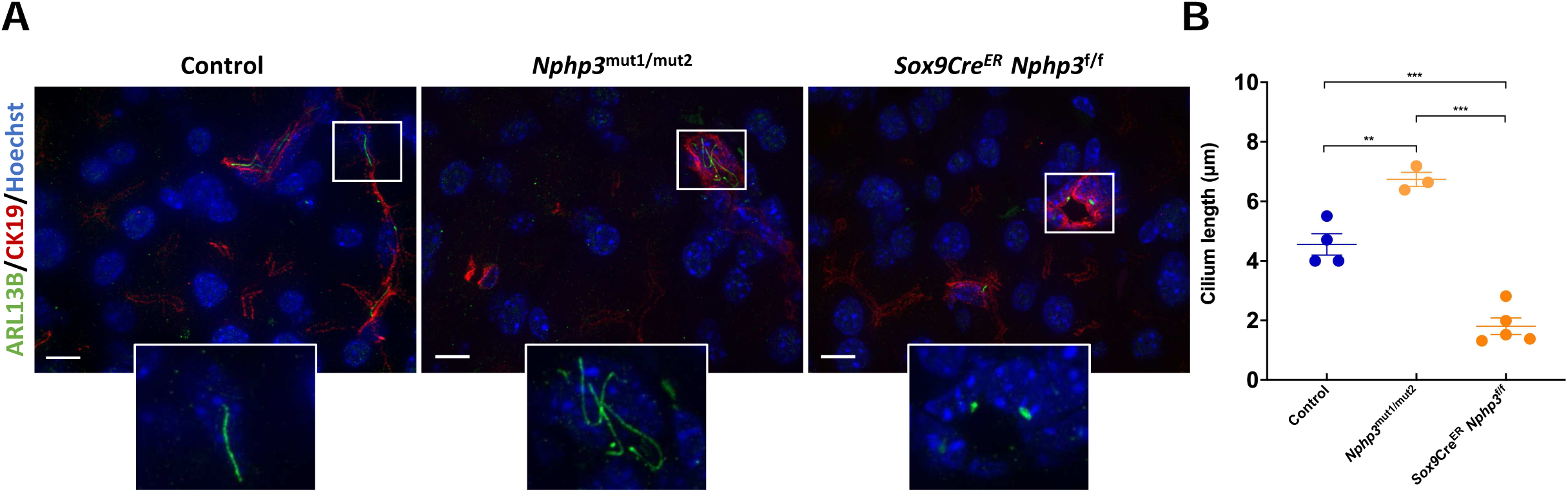
Cilium size is altered in *Nphp3* mouse models. **(A)** Immunolabeling of pancreas sections from Control, *Nphp3*^mut1/mut2^, and *Sox9Cre^ER^ Nphp3*^f/f^ mice at 8 months with ARL13B (for cilia) and CK19 (for ductal cells) antibodies, and Hoechst (nuclei, blue). Scale bar: 100 μm. **(B)** Quantification of the cilium length in the pancreas of Control, *Nphp3*^mut1/mut2^, and *Sox9Cre^ER^ Nphp3*^f/f^ mice. n=3-5 mice, and n=50 cilia counted per mouse.

### Ductal and centroacinar cells are differentially affected by patient-derived mutations and duct-specific inactivation of *Nphp3*

To further characterize ductal cells in our mouse models, we analyzed the expression of the transcription factor SOX9, which is present throughout the ductal tree, from the main pancreatic duct to the centroacinar cells located at the ends of the ducts in contact with the acinar cells^39^. Pancreatic inactivation of SOX9 during embryogenesis leads to failure in ductal cell differentiation, resulting in polycystic ducts devoid of primary cilia^40^.

SOX9 labeling initially revealed fewer SOX9-positive cells in *Nphp3*^mut1/mut2^ pancreas, compared to controls, in both 6- and 8-month-old mice (Figure 6A). Closer examination suggested a decrease in isolated SOX9-positive cells, likely corresponding to cells of the terminal ducts or centroacinar cells. Quantification confirmed an approximately 25–30% decrease in isolated SOX9-positive cells in the *Nphp3*^mut1/mut2^ pancreas (Figure 6B). In contrast, the width of the ducts was not significantly increased in *Nphp3*^mut1/mut2^ pancreata, although a non-significant trend toward larger ducts was observed at 8 months (Figure 6C). In the pancreas of *Sox9Cre^ER^ Nphp3*^f/f^ mice, we observed a striking, nearly threefold, reduction in isolated SOX9-positive cells, accompanied by an increase in the number of ducts, which appeared slightly dilated, in all size categories (small, medium and large) (Figure 6D–E). These results suggest that *NPHP3* mutations, whether hypomorphic or null, lead to a loss of SOX9 expression specifically in terminal ductal/centroacinar cells, or the absence of the latter, while SOX9 expression remains preserved in other ductal cells. They further indicate that pancreatic duct dilation occurs only with complete loss of NPHP3.

**Figure 6.**
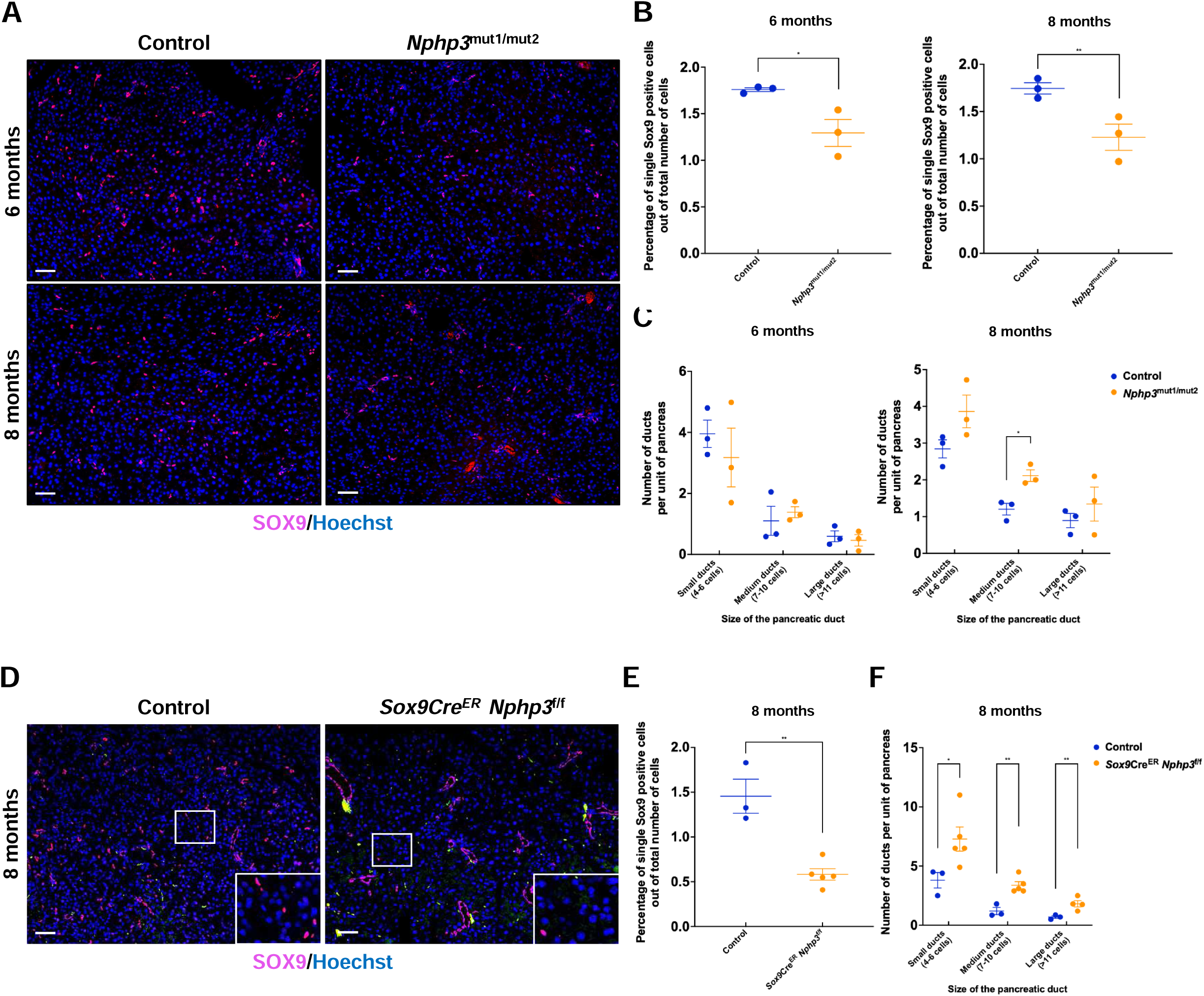
Ductal cells and pancreatic ducts are affected differently in Nphp3 mouse models. **(A)** Immunolabeling of pancreas sections from Control and *Nphp3*^mut1/mut2^ mice with SOX9 antibodies, and Hoechst. Scale bar: 50 μm. **(B)** Quantification of the percentage of isolated SOX9-positive cells out of total number of SOX9-positives cells in the pancreas of Control and *Nphp3*^mut1/mut2^ mice at 6 and 8 months. **(C)** Quantification of the number of ducts per unit of pancreas at 6 and 8 months. Ducts were classified according to their size (small, medium, large), based on their number of constituent cells. **(D)** Immunolabeling of pancreas sections from Control and *Sox9CreER Nphp3*^f/f^ mice with SOX9 antibodies, and Hoechst. Scale bar: 50 μm. **(E)** Quantification of the percentage of isolated SOX9-positive cells out of total number of SOX9-positive cells in the pancreas of Control and *Sox9CreER Nphp3*^f/f^ mice. **(F)** Quantification of the number of ducts per unit of pancreas.

### Acinar atrophy may result from apoptotic death

A notable feature of the *Nphp3* mouse models, also observed in HNF1B models^8, 23^, is the presence of acinar atrophy. To investigate the mechanisms underlying the progressive loss of acinar cells, we first analyzed the expression of MIST1, a transcription factor important for maintaining acinar cell identity^41^. We found a similar MIST1 expression in control and both *Nphp3* models, without any area exhibiting a significant reduction in MIST1 expression (Supplementary Figure 5A). This suggests that acinar identity was not affected by *NPHP3* mutations.

We next investigated whether acinar loss was associated with apoptotic cell death. In controls, TUNEL assay revealed only rare apoptotic events in acinar cells. In contrast, in mutant *Nphp3* mice, apoptotic acinar cells were occasionally detected, often appearing in clusters within certain pancreatic lobes, albeit with marked heterogeneity across the pancreas (Supplementary Figure 5B). These findings suggest that apoptosis contributes, at least in part, to the acinar atrophy observed in these models.

### Microcysts originating from secretory canaliculi are observed in the *Nphp3* mouse models

In certain areas of the pancreatic parenchyma of *Sox9Cre^ER^ Nphp3*^f/f^ mice, we observed histological abnormalities suggestive of large vacuoles within acinar cells (Figure 7A). To further characterize these structures, we performed immunolabeling for amylase (AMY), an enzyme produced by acinar cells, Mucin1 (MUC1), which is found on the apical surface of epithelial cells, and Zona Occludens 1 (ZO1), a tight junction protein. This analysis revealed the presence of microcyst-like structures that were MUC1-positive and ZO1-negative within the cytoplasm of acinar cells in the pancreas of *Sox9Cre^ER^ Nphp3*^f/f^ mice (Supplementary Figure 6A). These microcysts were often strongly positive for AMY, suggesting that amylase was concentrated within these microcysts and along their membranes (Figure 7B). Comparable structures were detected in the pancreas of the patient-derived model, though they appeared smaller in size (Figure 7B and Supplementary Figure 6A). The presence of the cysts was not spatially associated with other histopathological features (adipopancreatosis, or immune cell infiltration) observed in *Nphp3* models.

**Figure 7.**
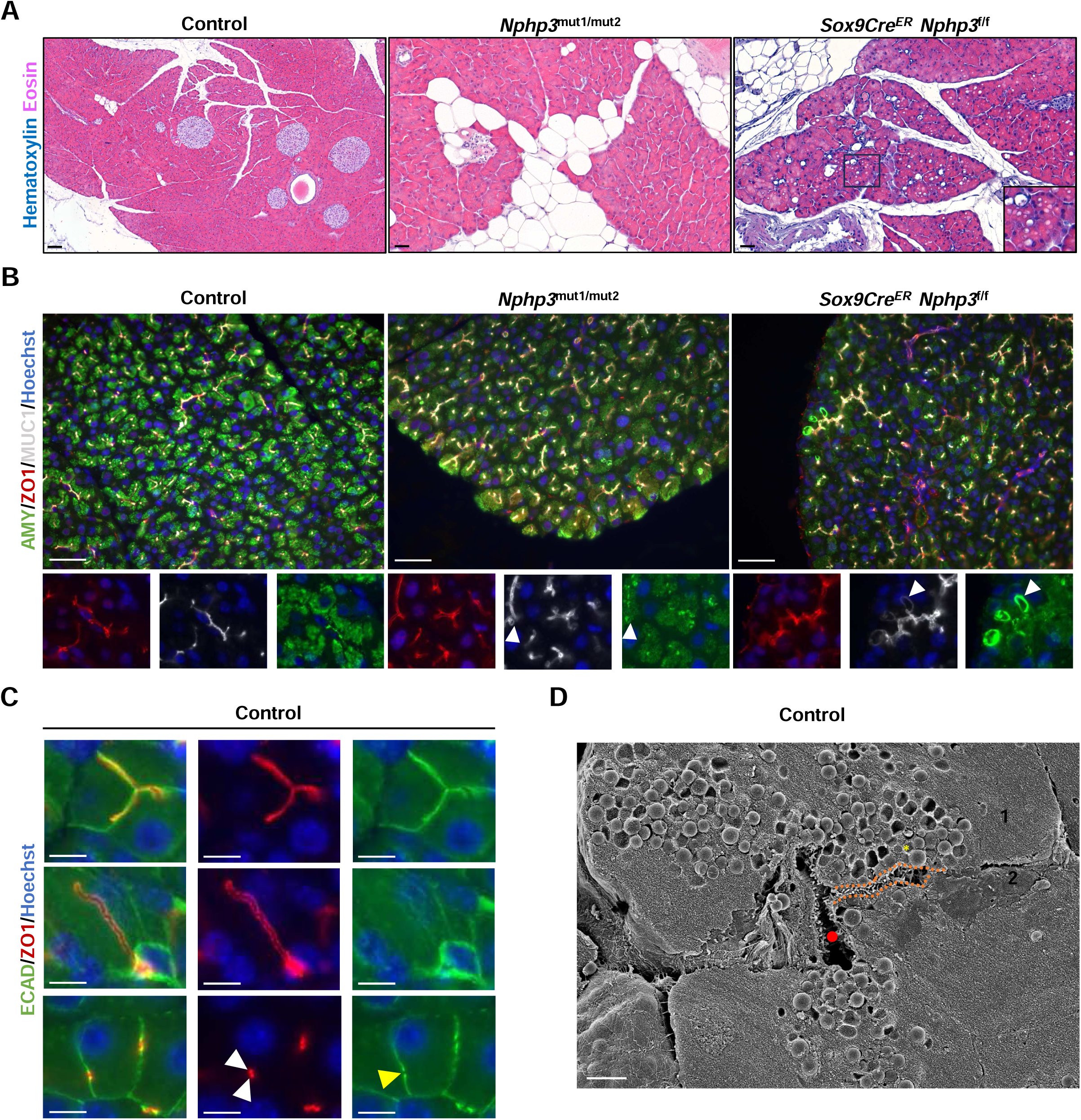
Acinar microcysts derived from secretory canaliculi are found in the pancreas of *Nphp3* mouse models. **(A)** Hematoxylin/Eosin staining of pancreas sections from Control, *Nphp3*^mut1/mut2^, and *Sox9CreER Nphp3*^f/f^ mice at 8 months. Vacuoles are observed in acinar cells of *Sox9CreER Nphp3*^f/f^ pancreata (inset). Scale bar: 100 μm. **(B)** Immunolabeling of pancreas sections from Control (3/3 mice showing no acinar microcysts), *Nphp3*^mut1/mut2^ (3/5 mice showing acinar microcysts in at least one pancreatic lobe), and *Sox9CreER Nphp3*^f/f^ (6/6 mice showing acinar microcysts in at least one pancreatic lobe) mice at 8 months with amylase (AMY), ZO1, and Mucin1 (MUC1) antibodies, and Hoechst. Scale bar: 50 μm. **(C)** Immunolabeling of pancreas sections from Control mice at 8 months with E-cadherin (ECAD, for the lateral pole of acinar cells) and ZO1 antibodies, and Hoechst (nuclei, blue). The upper images show ZO1 labeling superimposed on ECAD labeling along the lateral pole. The middle images show a secretory canaliculus extending across the cytoplasm of an acinar cell. The lower images show an interruption of the ECAD labeling at a precise location of the lateral pole (yellow arrowhead). At this location, the ZO1 labeling is characterized by a double dot (white arrowheads). Scale bar: 5 μm. **(D)** Scanning electron microscopy of a pancreas section from a Control mouse at 8 months. Red circle: central lumen. 1 and 2: two adjacent acinar cells. Orange dotted lines: intercellular secretory canaliculi with its microvilli. Yellow star: secretory granule. Scale bar: 2 μm.

To gain insight into these structures, we performed additional labeling with E-cadherin (ECAD), which is notably present at the lateral poles of acinar cells. In both control mice and *Nphp3* mouse models (Figure 7C and data not shown), two distinct patterns were redundantly observed at the lateral poles: either ZO1 and Ecad exhibited colocalization along a large portion of the lateral membrane (Figure 7C, upper part), or ZO1 appeared as a punctate signal present at the lateral pole (Figure 7C, lower part). In the latter case, ZO1 labeling was characterized by two closely spaced but separate dots, coinciding with a local interruption in ECAD expression at the same site. To complement these findings, we performed scanning electron microscopy, which revealed the presence of fine, microvillus-covered tubules within acinar cells, around which zymogen granules were concentrated (Figure 7D). Based on these observations, we propose that these structures correspond to intercellular secretory canaliculi located between adjacent acinar cells. This interpretation is further supported by former studies in the rat pancreas, which described identical structures using electron microscopy^42, 43^. In addition to intercellular secretory canaliculi, we also identified intraacinar secretory canaliculi, which are present within individual acinar cells (Figure 7C, middle part; Supplementary Figure 6B).

To further support this interpretation, we then performed 3D labeling on control pancreata. It confirmed the presence of both types of secretory canaliculi (Supplementary Figure 6C and Supplementary Movies 1-2). In *Nphp3* models, the microcysts described above were observed with increased frequency and size in the conditional knockout pancreas. These microcysts were located within the acinar cells and connected to their apical pole. Interestingly, acini containing microcysts generally lacked secretory canaliculi, indicating that the two structures were largely mutually exclusive, although in rare cases a cyst was found at the termination of a canaliculus (Supplementary Movies 3-4 and data not shown). 3D labeling also showed the presence of dilated ducts in regions of the *Sox9Cre^ER^ Nphp3^f/f^* pancreas where microcysts were not found. (Supplementary Movies 5). Taken together, these findings suggest that disruption of primary cilium function in ductal cells compromises the integrity of the network of secretory canaliculi and favors the development of acinar microcysts.

### Acinar microcysts are detected in other mouse ciliary models and in humans

To assess whether the presence of microcysts is a common feature of ciliary dysfunction, we examined additional ciliary models, namely mice with duct-specific inactivation of *Hnf1b*^8^ or *Intraflagellar Transport 88* (*Ift88*)^21^, which is essential for cilium formation, as well as a mouse model carrying an *HNF1B* mutation found in a patient^23^. Perilipin 1 labeling confirmed the presence of adipopancreatosis in these models, similar to our observations in the *Nphp3* models (Figure 8A). Subsequent labeling for Mucin1 revealed that microcysts were consistently present within the pancreatic parenchyma across all models (Figure 8B).

**Figure 8.**
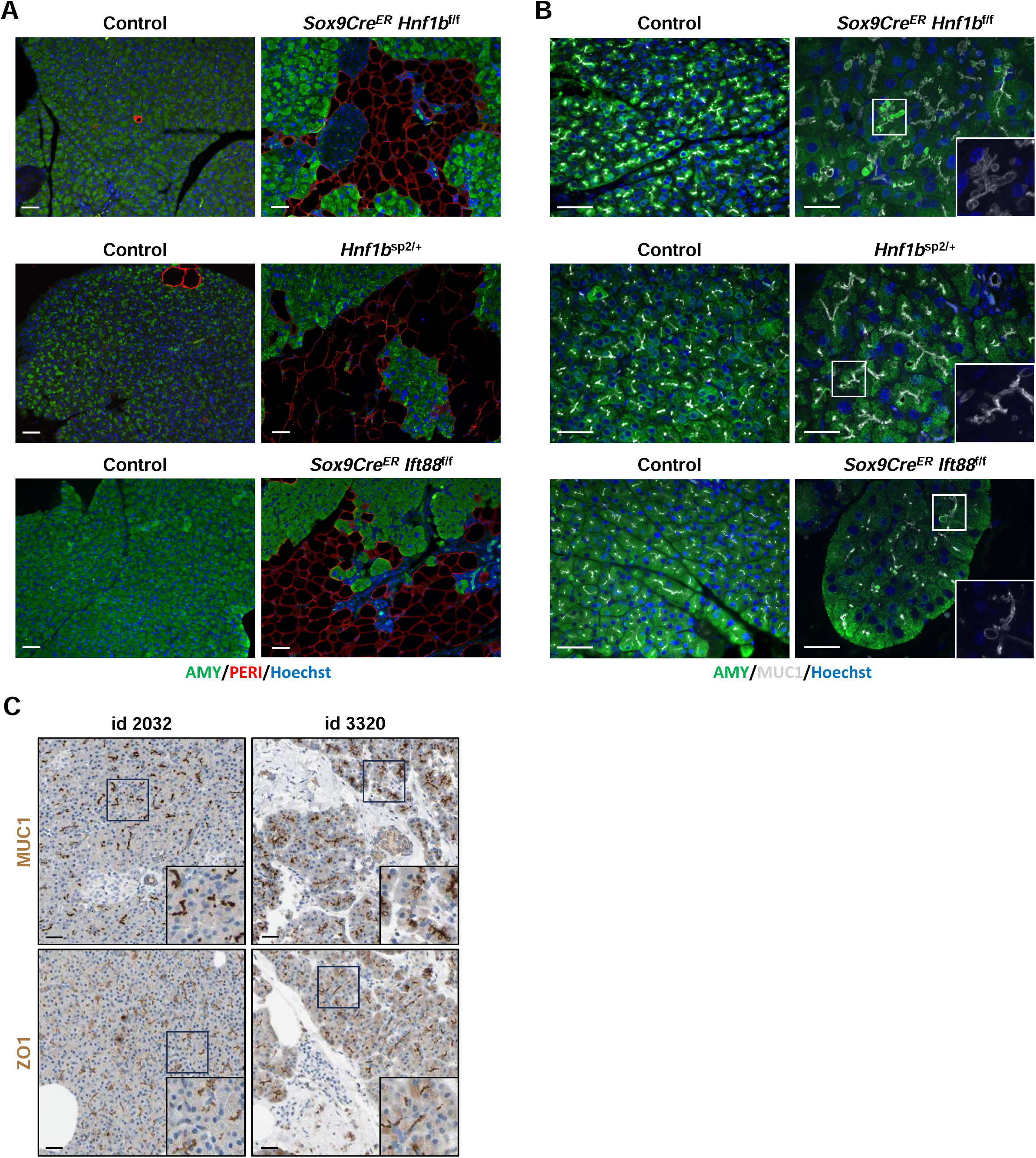
Acinar microcysts are present in additional ciliary mouse models and in human pancreas. **(A)** Immunolabeling of pancreas sections from Control (0/8 mice with adipopancreatosis), *Sox9CreER Hnf1b*^f/f^ (2/2 mice with adipopancreatosis), *Hnf1b*^sp2/+^ (3/3 mice with adipopancreatosis), and *Sox9CreER Ift88*^f/f^ (3/3 mice with adipopancreatosis), mice at respectively 2, 8 and 12 months with Perilipin 1 (PERI) antibody, and Hoechst. Scale bar: 50 μm. **(B)** Immunolabeling of pancreas sections from Control (0/8 mice with acinar microcysts), *Sox9CreER Hnf1b*^f/f^ (2/3 mice showing acinar microcysts in at least one pancreatic lobe), *Hnf1b*^sp2/+^ (2/2 mice with acinar microcysts in at least one pancreatic lobe), and *Sox9CreER Ift88*^f/f^ (2/3 mice with acinar microcysts in at least one pancreatic lobe) mice at respectively 2, 8 and 12 months with Mucin 1 (MUC1) antibody, and Hoechst. Scale bar: 50 μm. **(C)** Mucin1 and ZO1 staining of human pancreas sections from The Human Protein Atlas Resource (www.proteinatlas.org). Sample ID 2032 corresponds to a 32-year-old woman. Similar staining patterns were observed in pancreatic sections from 10 additional individuals. Sample ID 3320 corresponds to a 70-year-old woman. Medical data for this woman was not disclosed by The Human Protein Atlas Resource. The MUC1-positive microcysts observed in this pancreas were not found in 11 other pancreatic samples. Scale bar: 20 μm.

Building on these results in mice, we next investigated whether acinar microcysts might also occur in the human pancreas. To investigate this, we analyzed publicly available Mucin1 and ZO1 staining data from the Human Protein Atlas resource (HPA; proteinatlas.org). In human pancreatic tissue, Mucin1 staining was largely restricted to the center of the acini, with minimal extension toward the lateral poles (Figure 8C, top left). ZO1 staining exhibited a similar pattern (Figure 8C, bottom left). Interestingly, one pancreatic sample of the HPA resource displayed Mucin1-positive microcyst-like structures (Figure 8C, top right), similar to those observed in mouse models of ciliary dysfunction. These structures lacked ZO1 staining on a nearby section of the same pancreas (Figure 8C, bottom right), a situation closely resembling what is observed in the mouse pancreas. Unfortunately, information related to the pathophysiological status of the donor is not available. Nevertheless, our findings indicate that secretory canaliculi of variable length are detectable in rodent and human pancreas, and suggest that pancreatic disease can induce development of acinar microcysts in humans.

### Adipocytes in adipopancreatosis exhibit a transcriptional profile similar to white adipocytes and may derive from resident mesothelial fibroblasts

To characterize the pancreatic adipocytes at the molecular level, we microdissected adipocyte-enriched regions from the pancreas of an *Nphp3*^mut1/mut2^ mouse and compared their transcriptomic profile to reference transcriptomes of the three major adipocyte types (white, brown, and beige)^26^. The transcriptome of pancreatic adipocytes from *Nphp3* mutants most closely resembled that of white adipocytes (Figure 9A).

**Figure 9.**
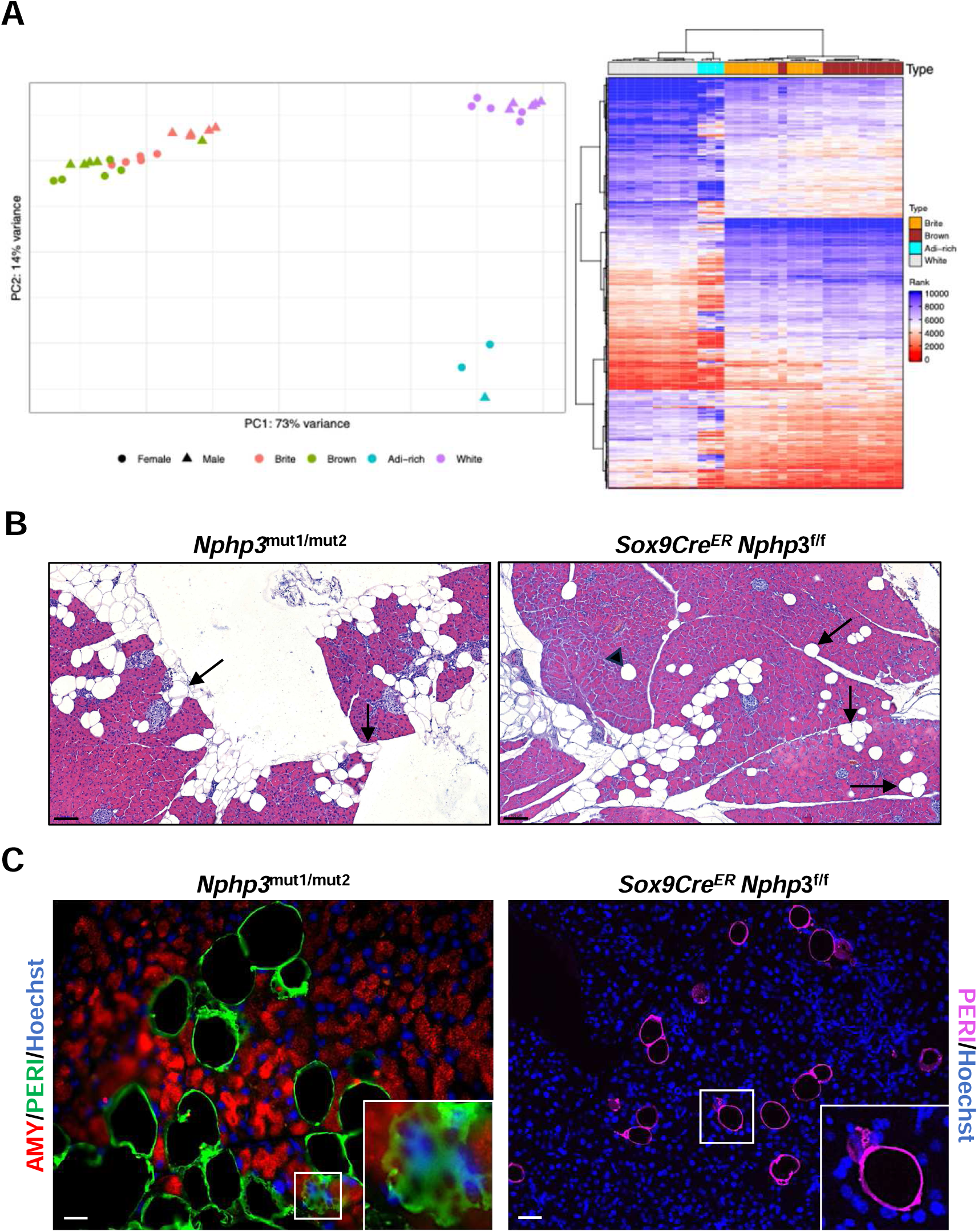
Principal component analysis (PCA) on rank matrix based on selection of 500 genes showing the greater variability in their ranking from brite, brown and white adipocytes and adipocyte-rich (Adi-rich) microdissected regions (left). Percentages indicate the proportion of variance explained by each component. **(B)** Hematoxylin/Eosin staining of pancreas sections from *Nphp3*^mut1/mut2^ and *Sox9CreER Nphp3*^f/f^ mice at 8 months. Arrow: groups of 2-3 adipocytes located at the periphery of pancreatic lobes/lobules. Arrowhead: isolated adipocyte found in the pancreatic parenchyma. Scale bar: 100 μm. **(C)** Immunolabeling of pancreas sections at 8 months from *Nphp3*^mut1/mut2^ mice with amylase (AMY) and Perilipin 1 (PERI) antibodies, and Hoechst (left picture) and from *Sox9CreER Nphp3*^f/f^ mice with Perilipin 1 (PERI) antibodies, and Hoechst (right picture). Scale bars: 10 μm (left) and 20 μm (right).

The morphology of the adipocytes in the *Nphp3* mutant models displayed interesting features. In regions where adipocyte accumulation began, the cells were primarily located at the periphery of the lobes, and to a lesser extent, closely apposed to the acini (Figure 9B). The lobes and acini are surrounded by mesothelial tissue comprising fibroblasts and it is known that fibroblasts can transdifferentiate into adipocytes under certain conditions^44, 45, 46^. It is therefore tempting to speculate that the adipocytes accumulating in the pancreas derive from resident mesothelial fibroblasts. Immunolabeling for Perilipin 1 revealed the presence of cells containing multiple cytoplasmic lipid droplets, a hallmark of preadipocytes. These preadipocytes were consistently located in close proximity to mature adipocytes, further supporting the possibility of *de novo* adipocyte formation (Figure 9C).

Taken together, our results indicate that the adipocytes accumulating in the pancreas display a white adipocyte profile and may derive from resident mesothelial-derived fibroblasts in contact with the ductal compartment via cellular extensions.

### Patients with *HNF1B* or *NPHP3* mutations have increased pancreatic fat content

The presence of adipopancreatosis in mouse models carrying ciliary mutations suggests that a similar situation occurs in the pancreas of patients with *HNF1B* or *NPHP3* mutations. As pancreatic tissue samples from these patients were not available, we assessed their PFC using Dixon-MRI, the reference method for quantitative PFC measurement, which separates water and lipid signals through in-phase and out-of-phase acquisitions^47^.

Ten patients with *HNF1B* mutations and three patients with *NPHP3* mutations (Figure 1A), along with 13 matched healthy controls, underwent Dixon-MRI. In some patients, no abnormalities of the pancreas were detected during diagnostic evaluations. PFC was significantly higher in patients with *HNF1B* (median: 7.4%; range: 3.6–19.3%) and *NPHP3* (median: 12.7%; range: 10.5–16.8%) mutations compared to their respective controls (*HNF1B* controls: median 2.1%, range: 1.7–4.2%, p = 0.0024; *NPHP3* controls: median 2.7%, range: 2.3–3.5%, p = 0.028) (Figure 10). These results support the hypothesis that, as in mouse models, patients with *HNF1B* or *NPHP3* mutations exhibit adipopancreatosis.

**Figure 10.**
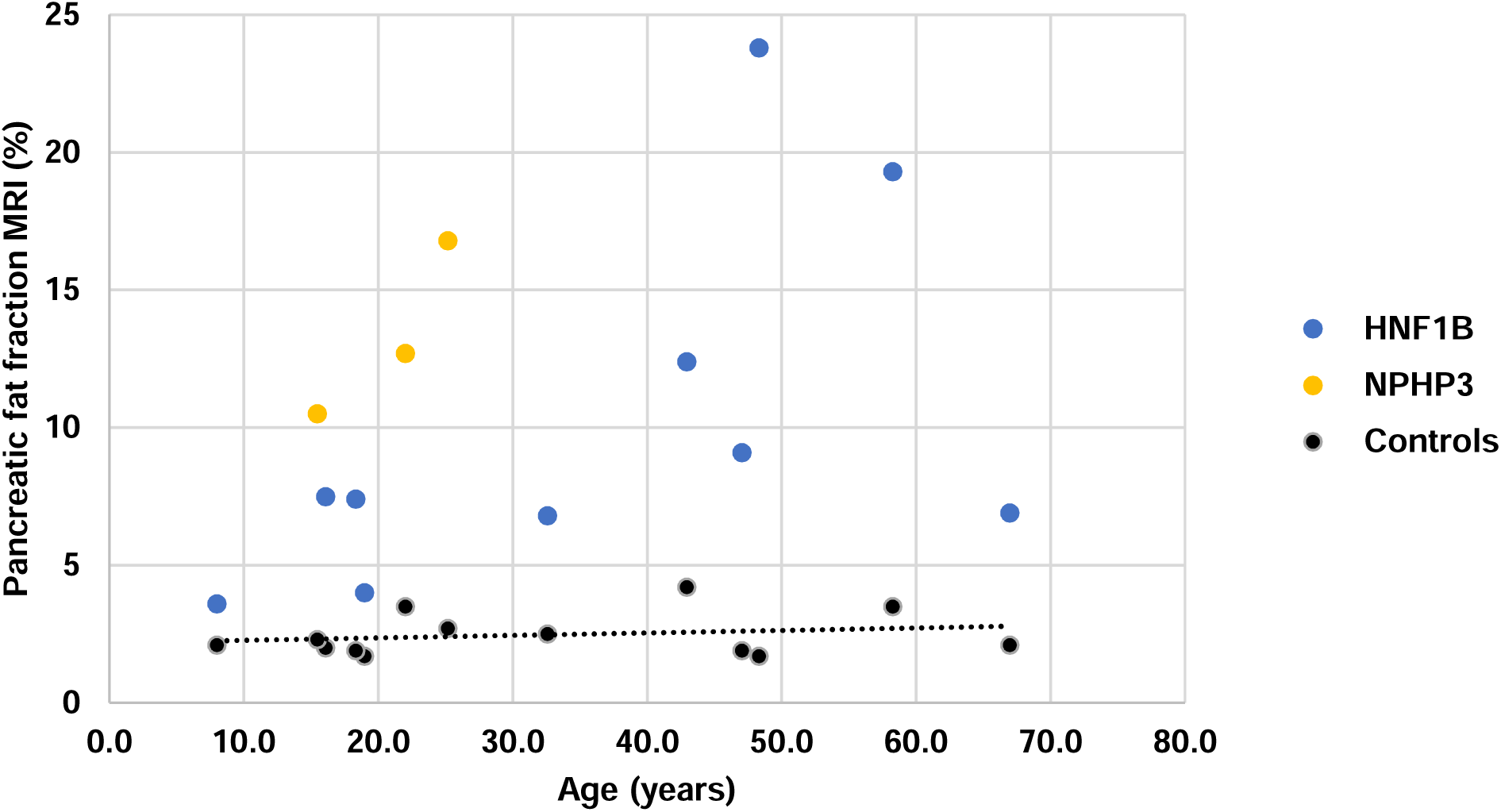
Pancreatic fat content as assessed by Dixon-MRI in healthy controls (black) and patients with *NPHP3* (yellow) and *HNF1B* (blue) mutations. Dashed line indicates the mean PFC in healthy controls matched for age, sex, and body mass index.

## Discussion

In this study, we identify *HNF1B* and *NPHP3* mutations as genetic drivers of a previously unrecognized pancreatic disease, which we propose to call ciliogenic pancreatopathy. Our findings expand the phenotypic spectrum of ciliopathies to include the pancreas and provide mechanistic insights into how ciliary dysfunction in a non-acinar compartment can lead to exocrine pancreatic pathology. The observation that *HNF1B* and *NPHP3* mutations, both known to cause renal ciliopathies, are associated with pancreatic abnormalities challenges the current understanding of organ involvement in ciliopathies. To date, pancreatic dysfunction associated with ciliary gene mutations, including in the NPHP3 gene^48, 49^, has been largely limited to descriptions of cystic anomalies. Our study is the first, to our knowledge, to link specific ciliary gene mutations to a reproducible, non-cystic pancreatic phenotype characterized by acinar tissue remodeling and fatty replacement, which we term adipopancreatosis. The presence of cysts in the reported studies^48, 49^ may be explained by the fact that the described mutations cause a more severe loss of NPHP3 activity, leading to death shortly after birth, unlike the patients in our cohort.

Clinically, this phenotype was supported by quantitative imaging using Dixon-MRI, which revealed significantly elevated PFC in both *HNF1B* and *NPHP3* mutation carriers compared to matched controls. Notably, in several of these patients, no other cause of pancreatic dysfunction could be identified, strengthening the causal link between *HNF1B* and *NPHP3* mutations and adipopancreatosis. This finding has immediate translational relevance: as with renal and hepatic involvement, pancreatic dysfunction may evolve silently in patients with ciliopathy-associated mutations, potentially leading to underdiagnosed exocrine insufficiency and associated nutritional complications. A potentially harmful clinical consequence relates to the coincidence of exocrine insufficiency and kidney disease. Exocrine insufficiency is known to cause steatorrhea, which in turn leads to increased intestinal absorption of oxalate, resulting in hyperoxaluria and promoting the formation of calcium oxalate stones, as well as oxalate nephropathy. The occurrence of this crystalline nephropathy could accelerate the decline in renal function in NPH patients, as observed in other kidney diseases, and may even contribute to an increased risk of graft rejection in NPH patients who have undergone kidney transplantation.

Mouse models were critical to unraveling the pathophysiology of ciliogenic pancreatopathy. Both the patient-specific *Nphp3^mut1/mut2^* model and the conditional *Sox9Cre^ER^ Nphp3*^f/f^ mice developed progressive adipopancreatosis and acinar atrophy, in the absence of significant fibrosis or overt acinar dedifferentiation. Importantly, these phenotypes emerged despite relatively mild inflammatory infiltration. The specific deletion of *Nphp3* in ductal cells was sufficient to recapitulate the adipopancreatosis phenotype, indicating that ciliary dysfunction in ductal, but not acinar, cells is a key initiating event. This aligns with the known ciliation pattern in the pancreas, where primary cilia are absent from acinar cells but present on ductal and centroacinar cells. Interestingly, ductal cilium length was either reduced or elongated depending on the model, suggesting that both hypomorphic and null mutations in *Nphp3* disrupt ciliary homeostasis, albeit through different mechanisms. It is well established that mutations in ciliary genes, as illustrated in particular by IFT88^50, 51^, can differentially affect ciliogenesis depending on whether they are hypomorphic or null, with distinct consequences on cilium formation, length, or function.

Our data also highlight the potential involvement of the stromal compartment, and specifically mesothelial-derived fibroblasts, in the emergence of pancreatic adipocytes. Immunolabeling revealed adjacent preadipocyte-like cells, and transcriptomic profiling showed that the adipocytes exhibited a white adipocyte–like phenotype. Together, these findings support a model in which resident fibroblasts, under the influence of local cues, transdifferentiate into adipocytes, thereby driving adipopancreatosis. Further reinforcing this hypothesis, *Snail1* knockout mice, lacking a key transcription factor for fibroblast identity, developed a similar phenotype. These results suggest that fibroblast fate maintenance is essential for preventing adipogenic conversion in the pancreas, and that ciliary signaling in ductal cells may contribute to this process.

Further, we identified another striking feature across multiple ciliary models, namely the presence of acinar microcysts originating from secretory canaliculi. These structures, positive for Mucin1, were localized between and within acinar cells and connected to their apical pole, consistent with previously described secretory canaliculi in rodent pancreas. Their increased size and frequency in *Nphp3* mutant mice, particularly in the conditional ductal knockout, suggest that ciliary dysfunction may secondarily affect acinar cell organization, possibly through altered duct-acinar signaling. The fact that similar microcyst-like structures were detected in human pancreatic samples from the HPA resource further supports their pathological relevance and raises the possibility that secretory canaliculi may be sensitive markers of early pancreatic dysfunction in ciliopathies.

Our findings have several important clinical implications. First, they expand the spectrum of ciliopathy-related organ involvement and highlight the need to systematically evaluate pancreatic function in patients with *HNF1B*, *NPHP3*, or other ciliary gene mutations, particularly in those presenting with unexplained gastrointestinal symptoms. Second, the presence of adipopancreatosis in these patients suggests a risk of exocrine pancreatic insufficiency, which may be subclinical but could negatively impact nutritional status and renal prognosis, particularly in transplant settings. Measurement of PFC using Dixon-MRI, a non-invasive and sensitive technique, could facilitate early detection of pancreatic involvement in ciliopathies. Future studies should also assess the prevalence of adipopancreatosis in other genetically confirmed ciliopathies, particularly those associated with mutations in genes encoding proteins that interact with NPHP3, such as NEK8 (NPHP9), and evaluate whether this process is reversible with cilia-targeted therapies.

Our study also has limitations. The cohort size remains limited for rare diseases, and pancreatic biopsy samples were not available from patients to confirm histological findings. However, the convergence of imaging, genetics, and mouse model data provides strong evidence for a true pathophysiological link. Further single-cell transcriptomic analyses of pancreatic stromal and ductal compartments may clarify the cellular trajectories leading to adipocyte emergence. Finally, our data support the notion that ciliogenic pancreatopathy is a progressive condition, with adipocyte accumulation and acinar atrophy worsening over time. Longitudinal follow-up of affected individuals will therefore be crucial to monitor disease evolution and guide early screening and timely therapeutic interventions.

## Accession number

The sequencing data associated with this study have been deposited in NCBI’s Gene Expression Omnibus (GEO) database and are accessible through GEO series accession numbers: GSE.

## Acknowledgments

We would like to thank Philippe Clapuyt for analyzing the pancreatic fat content by imaging, Luca Bertinetti and Yael Politi for usage and support of the SEM, and Angeline Fages, Raúl Peña, Nisha Limaye, Pascal Brouillard, and Raphaël Helaers for help. We thank the members of the histology and cell imaging facilities (S.F.R. Necker INSERM US24, CNRS UAR 3633 Paris, France) for technical assistance. This work was carried out between two centers that are members of ERNica and ERKNet.

## Ethics Statement

This study involves human participants and was approved by an Ethics Committee. The protocols were approved by the medical ethics review board of Cliniques universitaires Saint-Luc (CuSL) (PANC-GENET #2019/17DEC/568 and PANC-CILIA #2023/15MAR/135). This study involves animal subjects and was approved by an Ethics Committee. All experiments involving live mice were approved by the animal welfare committee of the University of Louvain Medical School (#2017/UCL/MD/020 and #2021/UCL/MD/054).

## Patient and public involvement

Patients or the public were not directly and actively involved in the design, or conduct, or reporting, or dissemination plans of our research.

## Abbreviations

ANKS6: ankyrin repeat and sterile alpha motif domain containing 6
CuSL: Cliniques universitaires Saint-Luc
DSSP: dictionary of secondary structure in proteins
HNF1B: hepatocyte nuclear factor 1 homeobox B
HNF6: hepatocyte nuclear factor 6
INVS: inversin
INVSc: inversin complex
MRI: magnetic resonance imaging
NEK8: NIMA-related kinase 8
NGS: next generation sequencing
NPHP3: nephrocystin 3
principal component analysis: PCA
protospacer adjacent motif: PAM
PDFF: proton density fat fraction
PFC: pancreatic fat content
PRSS1: serine protease 1
SPINK1: serine peptidase inhibitor Kazal type 1
TPR: tetratricopeptide repeats

