## supplemental figure for "Ciliogenic pancreatopathy reveals a link between ciliopathies and exocrine pancreatic disease"

### Supplementary Figure 1

**A**

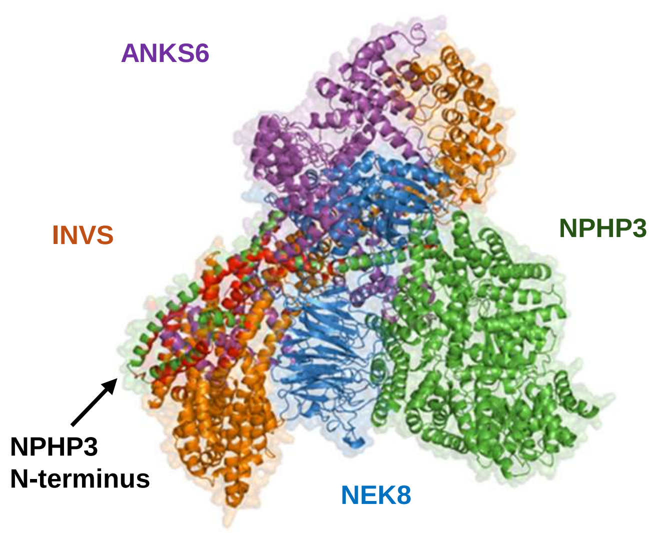


**INVSc, NPHP3 K144E**

**AlphaFold 3.0, model 0**

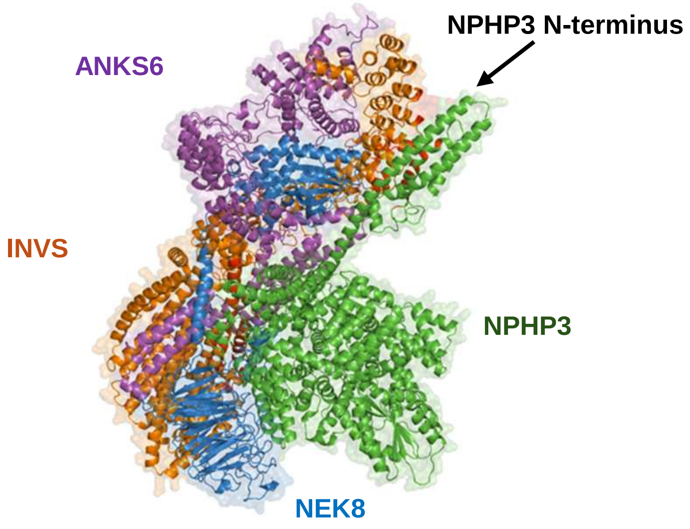


**INVSc, NPHP3 T307A**

**AlphaFold 3.0, model 0**

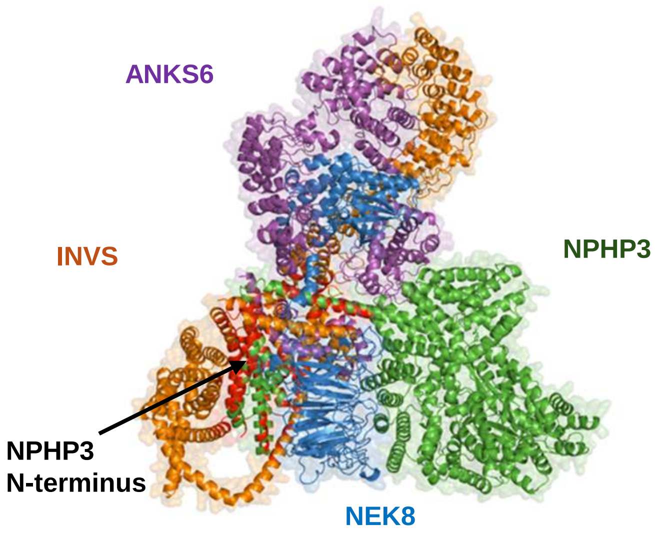


**INVSc, NPHP3 R1165Q**

**AlphaFold 3.0, model 1**

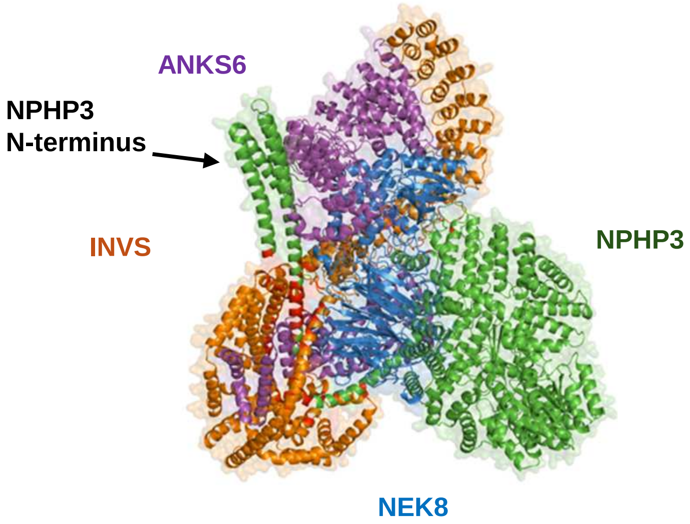


**INVSc, NPHP3 T1179R**

**AlphaFold 3.0, model 0**

**B**

| Mutant | Effect on topology of INVSc | Effect on contacts between NPHP3 and INVS | DynaMut prediction |
|---|---|---|---|
| **K144E** | INVS-NPHP3 N-terminus proximity conserved | Reduced number of contacts | Stabilizing + 0.87 kcal/mol |
| **T307A** | INVS-NPHP3 N-terminus proximity lost | Reduced number of contacts | Destabilizing - 1.0 kcal/mol |
| **R1165Q** | INVS-NPHP3 N-terminus proximity conserved | Similar number of contacts | Destabilizing - 0.95 kcal/mol |
| **T1179R** | INVS-NPHP3 N-terminus proximity lost | Reduced number of contacts | Destabilizing - 0.69 kcal/mol |

### Supplementary Figure 2

## A

| | mut1 | mut2 |
|---|---|---|
| Wild-type genomic sequence: | 5′-CAG AAG ATA CTT CGG GAA AAA GAG- 3′ | 5′-CTT CAG AAA ACT TAC CAG AAG ATA- 3′ |
| Wild-type protein sequence: | Q K I L R E K E | L Q K T Y Q K I |
| Mutated genomic sequence: | 5′-CAG AAG ATA CTT AGA GAG GAA GAG- 3′ | 5′-CTT CAG AAA -CT TAC AAG AAG ATA- 3′ |
| Mutated genomic sequence: | Q K I L R E **E** E | L Q K **L T R R Y** |

## B

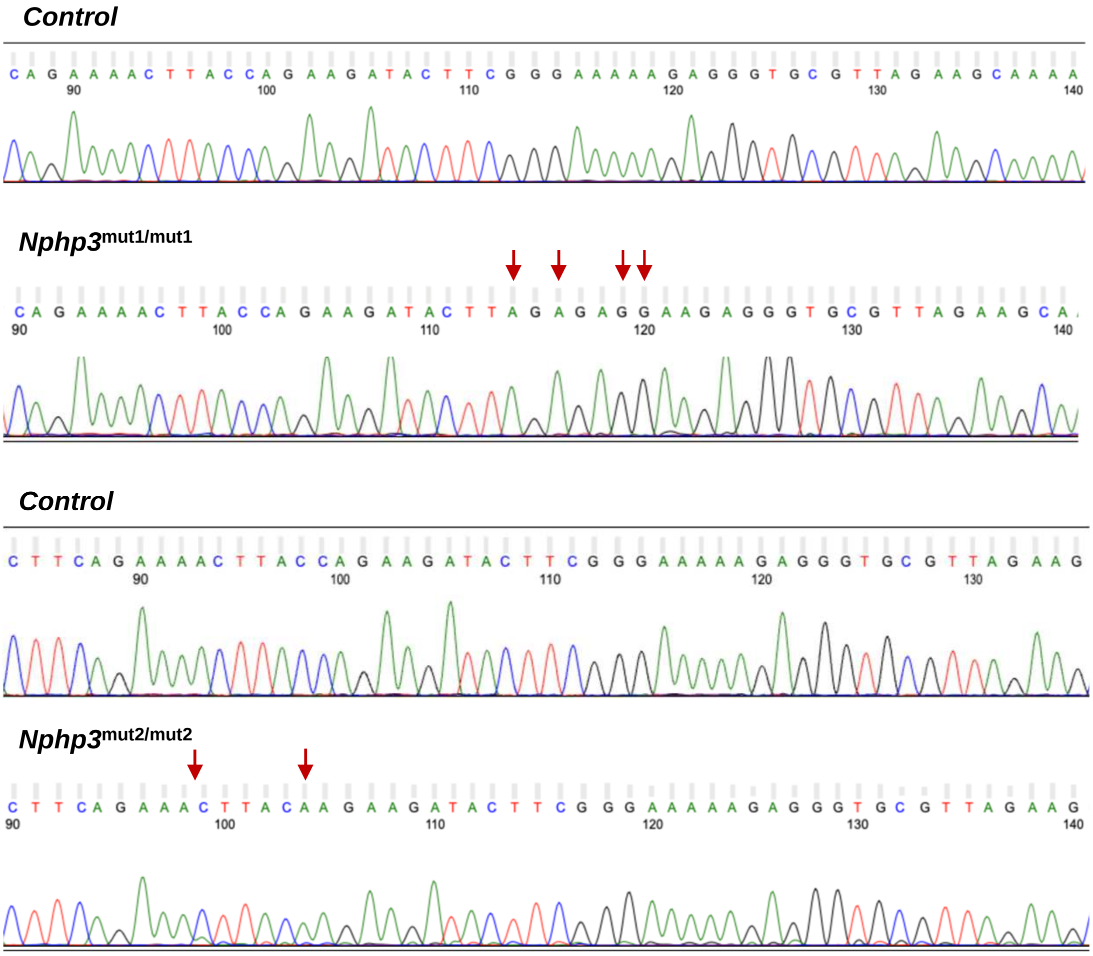


## C

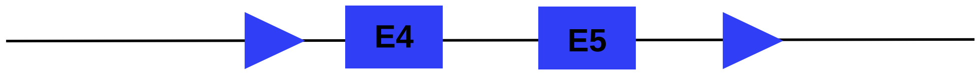

### Supplementary Figure 3

**A**

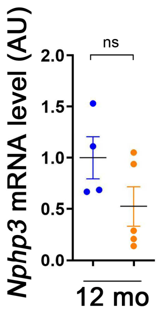


**B**

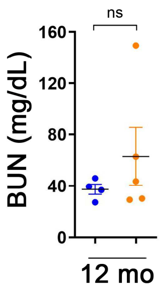


**C**

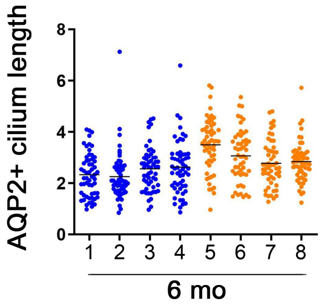

### Supplementary Figure 4

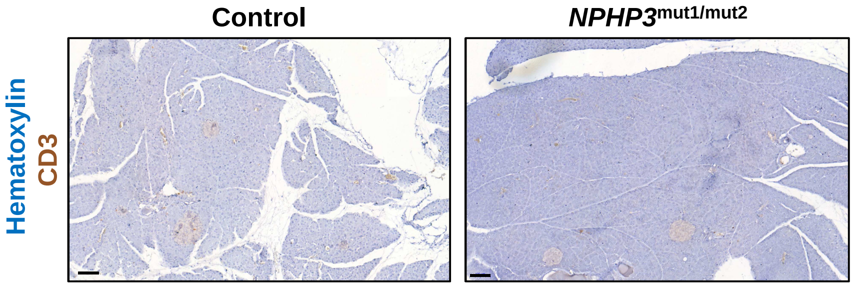

### Supplementary Figure 5

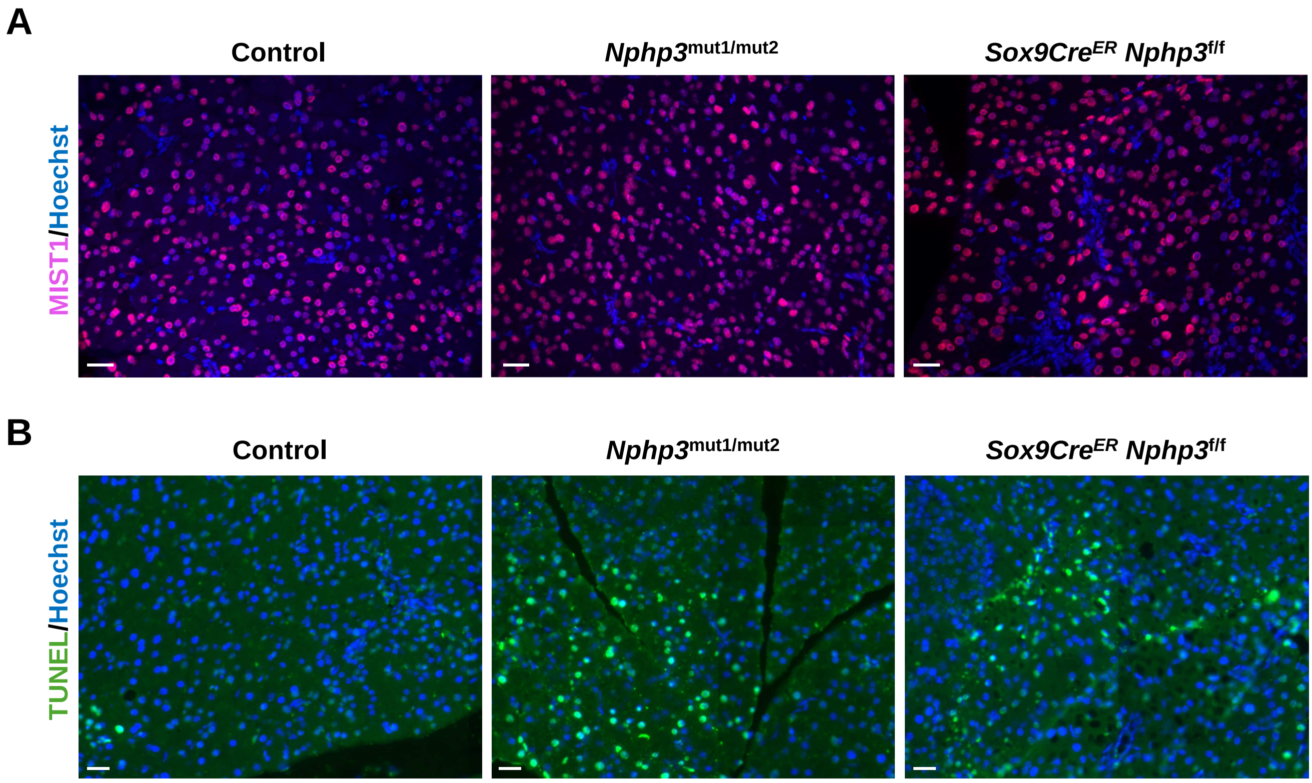


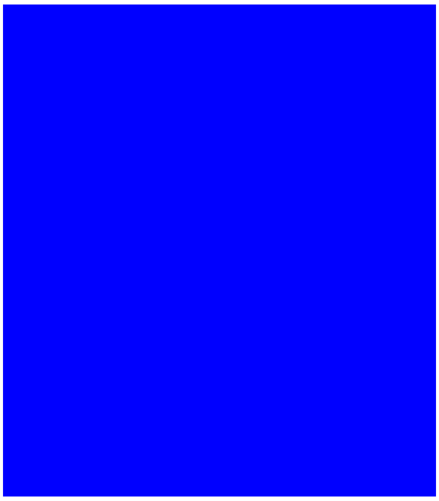

### Supplementary Figure 6

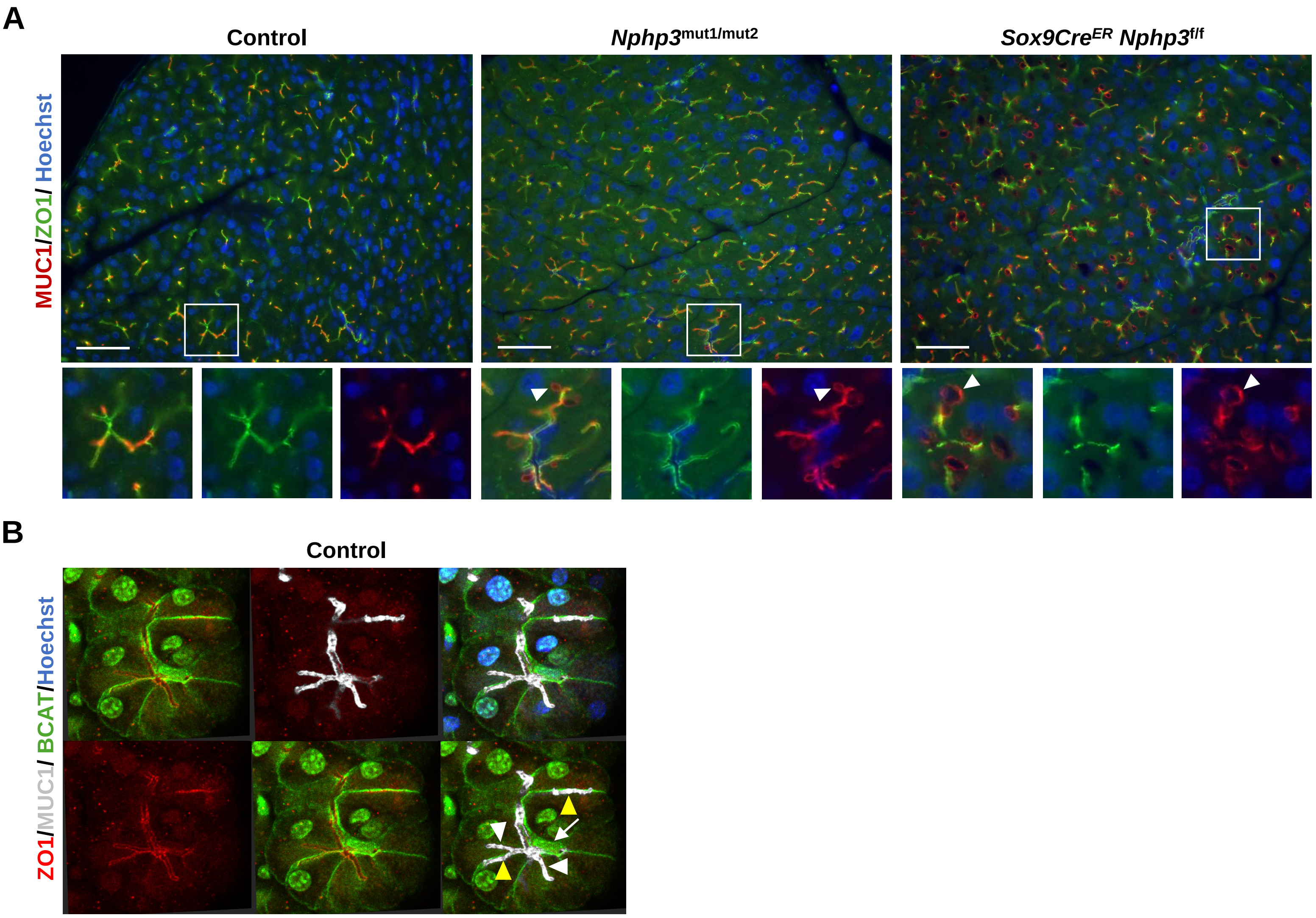

**Supplementary Figure legends**

**Supplementary Figure 1. (A)** Illustrative predictions of the INVS complex in the presence of the indicated NPHP3 mutants. ANKS6 is shown in purple, INVS in orange, NEK8 in dark blue, and NPHP3 in green. Contact residues between INVS and NPHP3 (distance < 6Ä) are shown in red. (**B)** Summary of *in silico* analysis of the effects of NPHP3 mutants on INVS complex.

**Supplementary Figure 2. Generation of mouse *Nphp3* models using CRISPR/Cas9 technology.** (**A**) Schematic showing CRISPR/Cas9-mediated introduction of the NPHP3_1 patient mutations into the mouse genome to generate the *Nphp3*$^{mut1/mut2}$ model. Wild-type (WT) and mutated mouse genomic sequences are shown (black), along with WT and mutated protein sequences (blue). Nucleotides affected by mut1 and mut2 mutations are shown in WT genome sequences (bold green. Nucleotide 655 for mut1 and nucleotide 631 for mut2 in NM_028721.3). Changes generated by CRISPR/Cas9 are shown in red in mutated genome sequences. For mut1, the nucleotide in bold red represents the nucleotide mutated in the patient and those in red represent the nucleotides mutated following the insertion of the single guide RNA sequence (silent mutations introduced to avoid further cleavage by Cas9 after introduction of the mutated sequence). The red dash indicates the deleted nucleotide within mut2. The altered amino acid within mut1 is shown in red. Mut2 introduces a frameshift that changes the amino acid composition, producing a protein with a predicted molecular weight of 16 kDa. (**B**) Chromatograms obtained by Sanger sequencing which show the WT, mut1, and mut2 (both in the homozygous state) DNA sequences. Red arrows indicate the location of mutated nucleotides introduced by CRISPR-Cas9-mediated genome editing. (**C**) Schematic illustrating the localization of LoxP sites introduced by CRISPR-Cas9 into the mouse *Nphp3* gene to generate the *Sox9*Cre$^{ER}$ *Nphp3*$^{f/f}$ model. E: exon.

**Supplementary Figure 3. *Nphp3*$^{mut1/mut2}$ mice have elongated cilia but no increase and plasma blood urea nitrogen. (A)** *Nphp3* mRNA content evaluated by qRT-PCR in controls and *Nphp3*$^{mut1/mut2}$ kidneys at 12 months. Each dot represents one individual mouse. Bars indicate the mean values. AU: arbitrary unit. **(B)** Plasma blood urea nitrogen (BUN) in controls and *Nphp3*$^{mut1/mut2}$ mice kidneys at 12 months. Each dot represents one individual mouse. Bars indicate the mean values. **(C)** Quantification of cilium length in AQP2 positive tubules (CD) in control and *Nphp3*$^{mut1/mut2}$ kidneys at 6 months. Each dot represents one individual primary cilium. Blue dot, Control; orange dot, *Nphp3*$^{mut1/mut2}$.

**Supplementary Figure 4. Lymphocytic infiltration is not present in the pancreatic parenchyma of *Nphp3*$^{mut1/mut2}$ mice.** Staining with a CD3 antibody and hematoxylin of pancreas sections from control and *Nphp3*$^{mut1/mut2}$ mice at 8 months. Scale bar: 100 µm. n=3-5.

**Supplementary Figure 5. Acinar atrophy is not the consequence of widespread apoptosis in *Nphp3* mouse models. (A)** Immunolabeling of pancreas sections from Control, *Nphp3*$^{mut1/mut2}$, and *Sox9Cre*$^{ER}$ *Nphp3*$^{f/f}$ mice at 8 months with MIST1 (for acinar cells) antibodies, and Hoechst (nuclei, blue). Scale bar: 50 µm. n= 3. **(B)** TUNEL assay performed on pancreas sections from Control, *Nphp3*$^{mut1/mut2}$, and *Sox9Cre*$^{ER}$ *Nphp3*$^{f/f}$ mice at 8 months. Nuclei were labeled with Hoechst. Scale bar: 20 µm. n= 3.

**Supplementary Figure 6. Acinar microcysts are found in the pancreas of *Nphp3* mouse models. (A)** Immunolabeling of pancreas sections from Control, *Nphp3*$^{mut1/mut2}$, and *Sox9CreER Nphp3*$^{f/f}$ mice at 8 months with ZO1 and Mucin1 (MUC1) antibodies, and Hoechst. Arrowhead: acinar microcyst. Scale bar: 50 µm. **(B)** 3D immunolabeling of pancreatic tissue from Control mice at 8 months with β-Catenin (for the cell membranes), Mucin1, and ZO1 antibodies, and Hoechst. White arrowhead, intraacinar secretory canaliculus; yellow arrowhead, intercellular canaliculus; white arrow, centroacinar cell nucleus.

**Supplementary Movie 1.** 3D projection of a whole-mount labeling of Mucin1 (grey), ZO1 (red), and β-Catenin (green) depicting the architecture of the exocrine compartment of a Control pancreas. Nuclei were labeled with DAPI (blue). Focus is made on intercalated ducts and connected secretory canaliculi branching in the acinus.

**Supplementary Movie 2.** Magnification of the 3D labeling shown in Supplementary Movie 1. Note the presence of intercellular canaliculi, positive for β-Catenin (green), and of intracellular canaliculi, negative for β-Catenin. Both types of canaliculi are positive for Mucin1 (grey) and ZO1(red).

**Supplementary Movie 3.** 3D projection of a whole-mount labeling of Mucin1 (grey), depicting the architecture of the exocrine compartment and the presence of microcysts in a $Sox9Cre^{ER}$ $Nphp3^{f/f}$ pancreas.

**Supplementary Movie 4.** 3D projection of a whole-mount labeling of Mucin1 (grey), ZO1 (red), and β-Catenin (green) depicting the architecture of the exocrine compartment and the presence of microcysts in a $Sox9Cre^{ER}$ $Nphp3^{f/f}$ pancreas. Nuclei were labeled with DAPI (blue).

**Supplementary Movie 5.** 3D projection of a whole-mount labeling of Mucin1 (grey) depicting the architecture of the pancreatic ductal tree in a $Sox9Cre^{ER}$ $Nphp3^{f/f}$ pancreas. Note the presence of a strongly enlarged duct in the periphery of the organ.

#### Supplementary Tables

**Supplementary Table 1.** NGS gene panel for clinical diagnosis of ciliopathies

| | | | | | | |
|---|---|---|---|---|---|---|
| ADAMTS9 | BBS5 | EYA1 | IQCB1 | NPHP1 | PTEN | TTC21B |
| ALG8 | BBS7 | FAN1 | LRP5 | NPHP3 | RPGRIP1L | TTC8 |
| ANKS6 | BBS9 | FH | LRP6 | NPHP4 | SDCCAG8 | UMOD |
| ARL6 | CEP164 | FLCN | LZTFL1 | OFD1 | SDHB | VHL |
| BBIP1 | CEP290 | GANAB | MAPKBP1 | PAX2 | SDHD | WDPCP |
| BBS1 | CEP83 | GLIS2 | MET | PKD1 | SEC63 | WDR19 |
| BBS10 | COL4A1 | HNF1B | MKKS | PKD2 | TMEM67 | XPNPEP3 |
| BBS12 | DCDC2 | IFT172 | MKS1 | PKHD1 | TRIM32 | ZNF423 |
| BBS2 | DNAJB11 | IFT27 | NEK8 | PMM2 | TSC1 | |
| BBS4 | DZIP1L | INVS | NOTCH2 | PRKCSH | TSC2 | |

**Supplementary Table 2. Relevant sequences for the generation of mouse models.**

**mNphp3-T132Lfs*13 single-strand oligonucleotide**

5’TGTAAAATGTGAACGAAAGGCCGTTTTTGTTAGGTGATAACAAGGTGAATCATTGCAG**GCACTTCAGAAACTT ACAAGAAGATACTTCGGGAAAAAGAGGGTGCGTTAGAAGCAAAATACCAAGCCATGGAGAGAGCGGTCACATT 3’**

**mNphp3-crRNA-T132Lfs*13**

**5’ CTTTTTCCCGAAGTATCTTC 3’**

**mNphp3-K140E single-strand oligonucleotide**

5’GGCCGTTTTTGTTAGGTGATAACAAGGTGAATCATTGCAG**GCACTTCAGAAAACTTACCAGAAGATACTTAGAGAGGAAGA GGGTGCGTTAGAAGCAAAATACCAAGCCATGGAGAGAGCGGTCACATTTGAACATGACAGAGACAG 3’**

**m-Nphp3-crRNA-K140E**

**5’ GAAGATACTTCGGGAAAAAG 3’**

NPHP3 patient-specific mouse model: nucleotides in bold correspond to exonic region. The arrows correspond to the desired nucleotide modifications del394A (pT132Lfs*13) and 418A>G (pK140E). The underlined nucleotides correspond to silent mutations introduced to destroy the PAM or the seed region.

**m-Nphp3 LoxP-Left-Ultramet**

5’AAACCACTACTTCAAGGCTTACTGTATTTACAAAAAGTGTATC**TTGATATGGGAACCCCT**GCTAGC**ataactt cgtata**ATGTATGC**tatacgaagttatATT**TGGCATATGTGGAGAGTTTTTAAACATTTGGTAACTTGGTTTTGA TGAAATTCCAAT3’

**m-Nphp3 LoxP-right-Ultramer**

5’GACCGCTCATGAGTGAAGAGCTCTGGCTGCTCTTGCAGAGGCCCCTGGTTTGTT**CCTCGG**GAATTC**ataactt cgtata**ATGTATGC**tatacgaagttatCACACATAAGCTGGCTC**ATGACTGGCTGTAATTCCTATTCCAGGGAAT ATTCCAGATTAG3’

**m-Nphp3-Left-crRNA:**

5’ TTGATATGGGAACCCCTATT3’

**mNphp3-right-crRNA :**

5’ GAGCCAGCTTATGTGTGCCG 3’

NPHP3 conditional knockout model: nucleotides in bold and underlined correspond to target sequence (20 nucleotides plus PAM). The nucleotides highlighted in yellow correspond to NheI and EcoRI restriction sites inserted into the intronic sequences close to the LoxP site. The nucleotides highlighted in purple correspond to the LoxP site sequence.

**Supplementary Table 3.** Primer pairs used for qRT-PCR.

| Gene name | Species | Forward Primer (5' to 3') | Reverse Primer (5' to 3') |
|---|---|---|---|
| ***Calb1*** | Mouse | TTGGCTCACGTCTTACCCACAGAA | ATGAAGCCGCTGTGGTCAGTATCA |
| ***Havcr1*** | Mouse | GAGAGTGACAGTGGTCTGTATTG | CCTTGTAGTTGTGGGTCTTCTT |
| ***Hprt*** | Mouse | GTTAAGCAGTACAGCCCCAAA | AGGGCATATCCAACAACAAACTT |
| ***Lcn2*** | Mouse | TCCTCAGGTACAGAGCTACAA | GCTCCTTGGTTCTTCCATACA |
| ***Nphp3*** | Mouse | ATGCCAAGTGACAGGGATAAA | CTGTGAACGATGGAGGTTAGAG |
| ***Ppia*** | Mouse | GGCTATAAGGGTTCCTCCTTTC | TTTCTCTCCGTAGATGGACCT |
| ***Rpl13*** | Mouse | GCTCCAAGCTCATCCTGTT | GGTGGCCAGCTTAAGTTCT |
| ***Umod*** | Mouse | TCAACAGAAGCGAGACGGTGTTCT | AGCACACTCATCCATGTCCTCACA |

**Supplementary Table 4.** Antibodies

| Antibodies | Origin | Dilution | Reference |
|---|---|---|---|
| Fluorescein Lotus Lectin | / | 1/200 | FL-1321 (Vector) |
| Barttin | Mouse | 1/200 | sc365161(A-3) (Sigma) |
| Uromodulin | Sheep | 1/400 | 8595-0054 (Biorad) |
| NCC | Rabbit | 1/200 | HPA028748 (Sigma) |
| Aquaporin 2 | Mouse | 1/200 | sc-9882 (Santa Cruz) |
| Calbindin | Chicken | 1/600 | NBP2-50028SS (Novus) |
| CD3 | Rabbit | 1/500 | ab16669 (Abcam) |
| F4/80 | Rabbit | 1/100 | D2S9R (BIOKE) |
| Amylase | Mouse | 1/500 | sc-46657 (Santa Cruz) |
| FABP4 | Rabbit | 1/300 | ab13979 (Abcam) |
| Perilipin 1 | Rabbit | 1/100 | PA572921 (Thermofisher) |
| Sox9 | Rabbit | 1/500 | Lab-made |
| GFP | Goat | 1/250 | ab6673 (Abcam) |
| ARL13B | Rabbit | 1/800 | 17711-1-AP (Sanbio) |
| Cytokeratin 19 | Rat | 1/500 | MABT913 (Merck) |
| ZO1 | Rabbit | 1/400 | 61-7300 (Thermofisher) |
| Mucin1 | Armenian hamster | 1/300 | m5a-11202 (Thermofisher) |
| E-Cadherin | Mouse | 1/200 | 610182 (BD Biosciences) |

| | | | |
|---|---|---|---|
| Vimentin | Rabbit | 1/300 | 5741 (Cell Signaling) |
| Mist1 | Rabbit | 1/100 | 14896 (Cell Signaling) |
